# UltimateSynth: MRI Physics for Pan-Contrast AI

**DOI:** 10.1101/2024.12.05.627056

**Authors:** Rhea Adams, Khoi Minh Huynh, Walter Zhao, Siyuan Hu, Wenjiao Lyu, Sahar Ahmad, Dan Ma, Pew-Thian Yap

## Abstract

Magnetic resonance imaging (MRI) is commonly used in healthcare for its ability to generate diverse tissue contrasts without ionizing radiation. However, this flexibility complicates downstream analysis, as computational tools are often tailored to specific types of MRI and lack generalizability across the full spectrum of scans used in healthcare. Here, we introduce a versatile framework for the development and validation of AI models that can robustly process and analyze the full spectrum of scans achievable with MRI, enabling model deployment across scanner models, scan sequences, and age groups. Core to our framework is UltimateSynth, a technology that combines tissue physiology and MR physics in synthesizing realistic images across a comprehensive range of meaningful contrasts. This pan-contrast capability bolsters the AI development life cycle through efficient data labeling, generalizable model training, and thorough performance benchmarking. We showcase the effectiveness of UltimateSynth by training an off-the-shelf U-Net to generalize anatomical segmentation across any MR contrast. The U-Net yields highly robust tissue volume estimates, with variability under 4% across 150,000 unique-contrast images, 3.8% across 2,000+ low-field 0.3T scans, and 3.5% across 8,000+ images spanning the human lifespan from ages 0 to 100.

---

Magnetic resonance imaging (MRI) offers a myriad of soft-tissue contrasts that are useful in physics, biology, and medicine. By adjusting scan parameters or selecting different scan sequences, such as T1-weighted, T2-weighted, and diffusion-weighted scans, specific tissue properties, such as water content, fat, or blood flow, can be highlighted, facilitating the diagnosis of conditions like tumors, inflammation, bleeding, and other pathologies. However, contrast variations arising from differences in imaging equipment, sequences, or site-specific protocols can cause inconsistent interpretation, diagnostic errors, and reduced reproducibility^1–6^. These variations also pose challenges in automated analysis—MRI computational tools often struggle to extract true tissue properties when faced with varying contrast. In large-scale datasets, such contrast variability further complicates data integration and analysis from multiple sources, reducing their collective clinical and research value.

AI networks for MRI often lack generalizability beyond their training data^7, 8^. This poses challenges in distinguishing actual tissue characteristics from contrast variations. Insufficient consideration of contrast deviations throughout the AI development life cycle—including data annotation, network learning, and model deployment—limits AI accuracy, reliability, and adaptability. Given the practical impossibility of gathering *in vivo* training images encompassing all conceivable contrasts, many contemporary approaches to improve MR image generalization revolve around synthesizing contrasts for data augmentation during network training.

Early MR physics-based simulation methods, such as SIMRI^9^ and MRiLab^10^, employed forward models to emulate realistic MR signal behavior under varying scanner settings and pulse sequences. However, these approaches are computationally intensive, as each component of the acquisition process—such as radiofrequency (RF) excitation, gradient application, and signal readout—must be individually simulated to generate the resulting images. To improve computational efficiency, closed-form MR signal equations can be used to simplify the simulation process. For example, static signal models corresponding to predefined sequences, such as MPRAGE, FLASH, and T2-SPACE, have been adopted to synthesize standard-looking images (e.g., SyntheticMR^11^) or to train segmentation models (e.g., PhysSeg^12^ and PSACNN^13^). These physically grounded simulators have laid a foundation for generating high-fidelity synthetic data. A key limitation of these methods, however, is their focus on mimicking images derived from existing MR sequences, rather than exploring the broader diversity of MR image contrasts— many of which may not align with any predefined sequence structure or scanner configuration. This restriction in simulation flexibility presents challenges for developing generalizable AI models, especially in scenarios involving novel or atypical images.

Alternatively, MR images can be synthesized without relying on physics-based or physiological modeling, using either domain randomization or deep generative models. Domain randomization^14^ assigns randomized, label-specific intensities to anatomical structures, producing atypical contrasts unconstrained by MR physics or tissue properties. This eliminates the need for paired multi-contrast data and improves generalizability without requiring domain adaptation. Segmentation networks trained entirely on such synthetic contrasts—such as SynthSeg^15^ and SynthSeg+^16^, which are inspired by Bayesian segmentation^17, 18^—have shown robust performance across a wide range of unseen MRI contrasts, resolutions, and scanner vendors. Deep generative models—including generative adversarial networks (GANs), variational autoencoders (VAEs), normalizing flows, and diffusion models—can synthesize anatomically plausible MR images by learning directly from empirical data^19^. However, these models are typically constrained by the distribution of their training data and often struggle to generalize to out-of-distribution scenarios. While domain randomization helps overcome this limitation, current implementations often neglect MR physics and natural tissue transitions, leading to oversimplified boundaries and potential segmentation errors.

Importantly, the use of synthetic contrasts has so far been largely confined to model training, overlooking their broader potential across the AI development pipeline. Particularly, their application in data labeling and performance benchmarking is conspicuously overlooked, despite the undeniable significance of these steps in AI development. Together, these limitations constrain the potential of synthetic contrast methods in realizing truly *pan-contrast* AI models—models that are robust to the full spectrum of MRI contrast variations.

Here, we introduce UltimateSynth—**U**ltrawide **L**andscape of **T**issue-faithful **I**mages for **M**ulticontrast **A**nnotation, **T**raining, and **E**valuation via Synthesis—a method built from the ground up based on the fundamental principles of MR physics to generate any desired MR image contrast on demand, with the ambitious goal of enabling pan-contrast generalizability for AI networks (Figure 1). Rather than relying on MR sequence-dependent models, UltimateSynth utilizes classic spin dynamics equations to produce a full spectrum of image contrasts across the complete range of tissue properties and scan parameters. This encompasses typical image contrasts from clinical scans, extreme contrasts attainable only through aggressive scanner settings, and suboptimal contrasts that may occur during routine acquisitions. Common contrasts such as T1-weighted, T2-weighted, and FLAIR represent only a small fraction of what can be generated with UltimateSynth. Additionally, UltimateSynth can synthesize contrasts beyond the physical limits of MRI scanners and sequences, equipping AI models to handle unforeseen imaging variations. The authenticity of the contrasts generated with UltimateSynth is a direct result of explicitly integrating intrinsic tissue properties, quantified through nuclear spin relaxation, and extrinsic imaging factors, like the static magnetic field, radiofrequency pulse, and acquisition timings. UltimateSynth’s ability to generate diverse and genuine images enables training AI models compatible with any MRI scan.

**Fig. 1.**
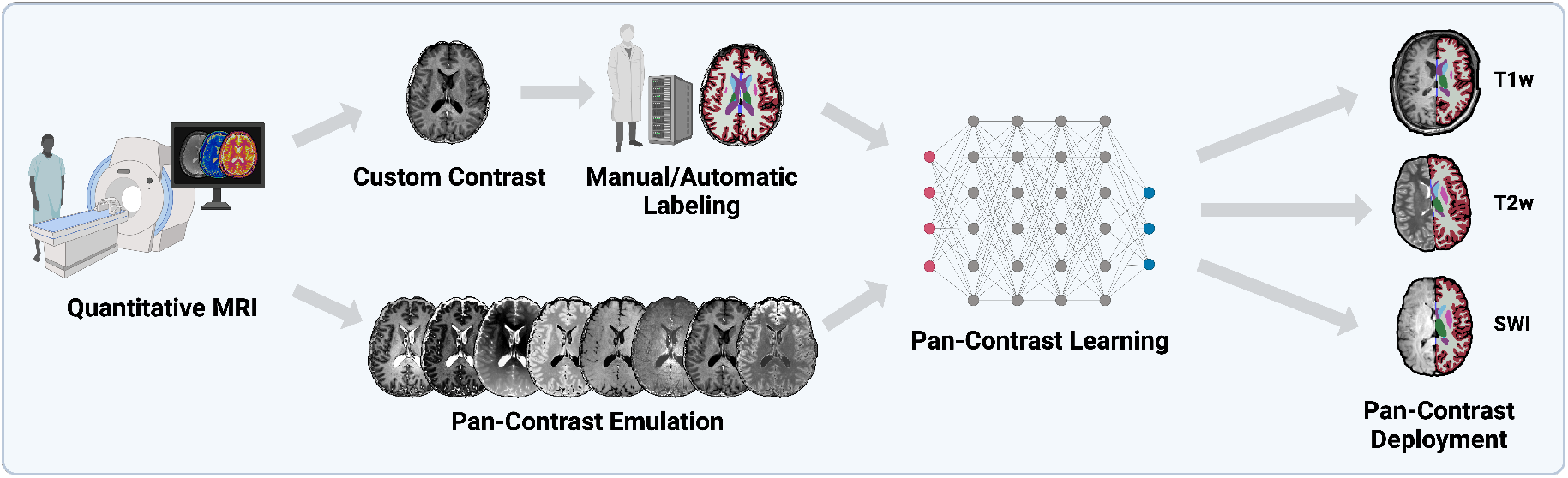
Pan-contrast AI via efficient annotation. UltimateSynth facilitates the learning of pan-contrast AI models by using data labeled only with as few as one contrast. It generates contrasts customized to software tools or annotator preferences, offering wide-ranging versatility in data labeling. UltimateSynth substantially reduces data acquisition and labeling efforts while boosting model output consistency by employing a diverse range of label-consistent contrasts during model training. In addition, UltimateSynth allows for the explicit validation of how well models handle the full range of MRI contrasts or any specific MRI contrast—a critical feature missing in existing MRI AI frameworks.

Importantly, UltimateSynth’s pan-contrast capability bolsters all three key stages of the AI development life cycle: Labeling, Emulation, and Deployment (LED). UltimateSynth offers customizable contrasts for efficient data labeling (Figure 2b), facilitates widely generalizable model training to avoid costly retraining (Figure 2c), and provides a platform for exhaustive performance benchmarking (Figure 2d):

**Fig. 2.**
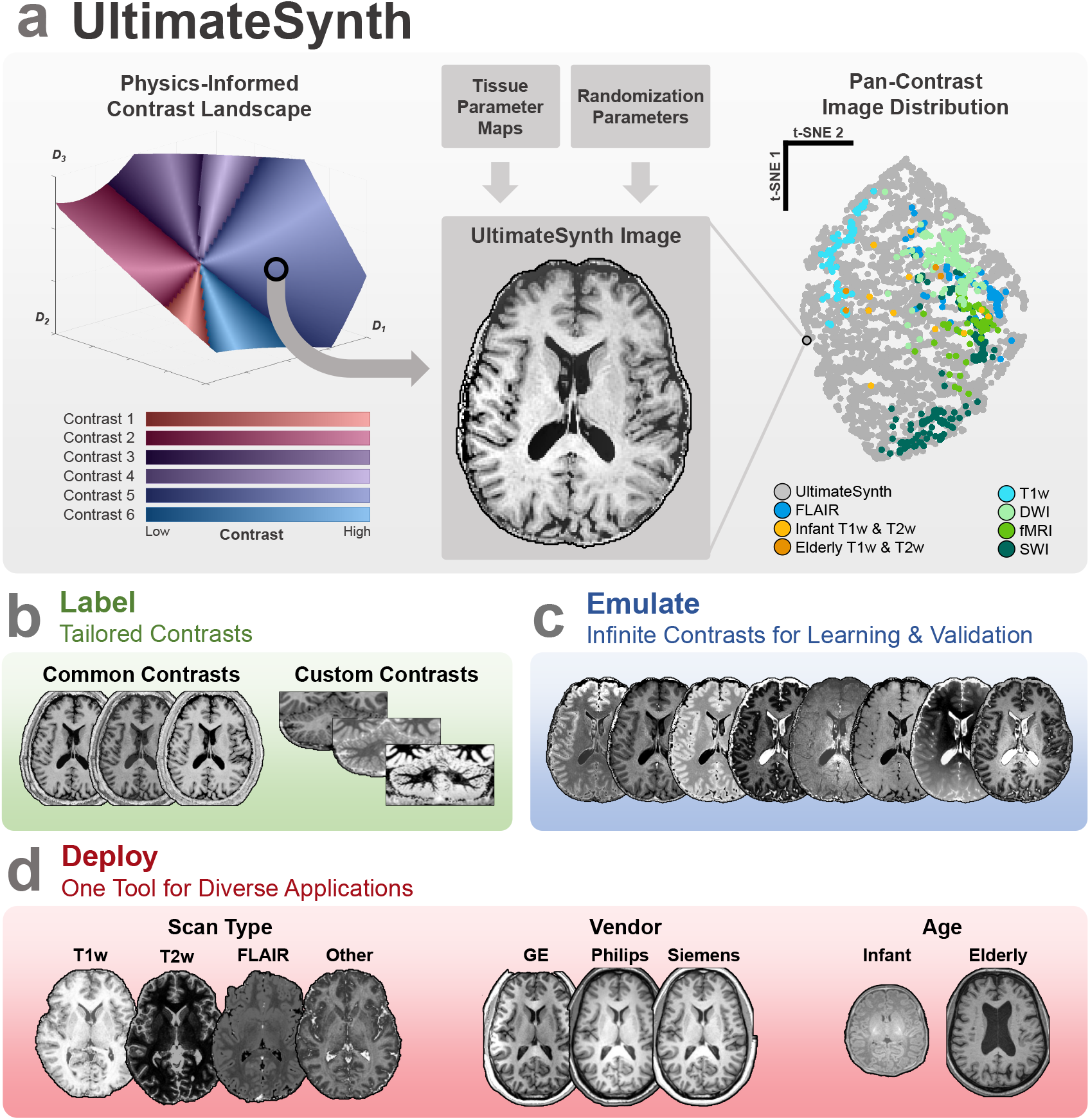
UltimateSynth and the AI LED life cycle. **a**, UltimateSynth uses MR physics to generate a contrast landscape based on diverse tissue and scan parameters. UltimateSynth images are produced by sampling from this landscape in combination with tissue parameter maps and random perturbations. The t-SNE^20^ plot of contrast radiomic features shows that UltimateSynth images fully encompass both common and uncommon MR images, highlighting their broad coverage. **b**, UltimateSynth generates tailored contrasts for automatic labeling or enhanced contrasts for easier manual delineation. **c**, UltimateSynth generates realistic and diverse images from a anatomical instance for pan-contrast model learning. **d**, UltimateSynth allows pan-contrast evaluation to ensure model generalizability across different contrasts, vendors, and ages.

## 1. Labeling

The first step in AI development involves annotating a training dataset, either manually or using automated tools like FreeSurfer^21^. However, conventional approaches often label only one image per anatomical instrance, necessitating considerable time to label large volumes of data for training. UltimateSynth substantially improves labeling efficiency by generating a wide spectrum of contrasts from the same anatomical instance.

This allows for the augmentation of training data and improves reliability by enforcing labeling consistency across different contrasts. UltimateSynth also enhances manual annotation by improving the visibility of structures with poor contrast, like the cerebellum, using customized image contrasts.

### 2. Emulation

The next stage of AI development involves training network models with labeled data. UltimateSynth substantially broadens training data by generating common, uncommon, and even practically unachievable contrasts beyond those used for labeling, enabling pan-contrast model training and maximizing the value of labeled data. Unlike methods that produce arbitrary contrasts, UltimateSynth faithfully emulates realistic MR contrasts grounded in physics and physiology.

### 3. Deployment

Before deployment, models must be thoroughly evaluated across various image contrasts. By simulating diverse and realistic images, UltimateSynth offers a platform for stress-testing AI models and quantifying uncertainty, ensuring model effectiveness across diverse datasets, vendors, and age groups.

We demonstrate with independent datasets spanning across scanners, acquisition protocols, and ages that a single U-Net^22^ segmentation model trained with UltimateSynth, using only a modest number of acquired images, yields unprecedented generalizability and consistency across all cases. Notably, when stress-tested with more than 150,000 unique MR contrasts, the model yields high consistency in volumetric quantification with exceptionally low mean volumetric variability of 3.35%, a nearly 5-fold improvement over the state-of-the-art SynthSeg^15^. We further evaluated the U-Net on external datasets to assess intra-subject repeatability, cross-lifespan repeatability, and low-resolution generalization, using 10,975 images across seven resolutions and six contrast types.

## RESULTS

### Realistic Pan-Contrast Synthesis

UltimateSynth simulates a comprehensive dictionary of signal magnetizations that reflect the combined effects of intrinsic tissue parameters (such as T1 and T2 relaxation times and proton density) and extrinsic acquisition parameters that determine contrast mechanisms (such as initial magnetization, pulse timings, and radiofrequency excitations). The simulation models classical nuclear spin dynamics without relying on anatomical or scan-specific assumptions. The UltimateSynth dictionary is then projected to a multidimensional subspace using singular value decomposition (SVD)^23^, yielding a physics-informed contrast landscape. Each coordinate in the subspace is associated with an eigenvector of the dictionary, and each element of the eigenvector corresponds to a unique combination of tissue and acquisition parameters. In Figure 2a, we visualize this subspace using the first three primary eigenvectors with the landscape colored according to different relative tissue contrasts for brain MRI (see Contrast Classes). With additional information from tissue property maps, any desired MR contrast can be generated for a target anatomy. Repeating this process across different subspace coordinates yields a diverse collection of unique images encompassing the full range of MRI scan types, as illustrated by the pan-contrast image distribution in Figure 2a.

In this work, we used tissue parameter combinations from *in vivo* human brain maps to generate diverse image contrasts—spanning common, uncommon, and unachievable contrasts— across high- and low-end scanners, incorporating spatial inhomogeneities from variable scanning conditions and random perturbations. Figure 3a illustrates that 3,200 images produced by UltimateSynth from a single subject fully encompass the diversity found in a set of 575 independent MR images. UltimateSynth images match the contrast variations seen across multiple scan types from external datasets (Figure 3b). Figure 3c shows that UltimateSynth can produce infinitely more images with drastically fewer subjects, sites, vendors, or ages needed than conventional MRI. We demonstrate in Movie S1 the pan-contrast diversity of UltimateSynth for one subject. Leveraging the infinite range of possible UltimateSynth contrasts reflecting underlying tissue parameters, we aim to improve the entire AI development LED life cycle (Figures 2b–d).

**Fig. 3.**
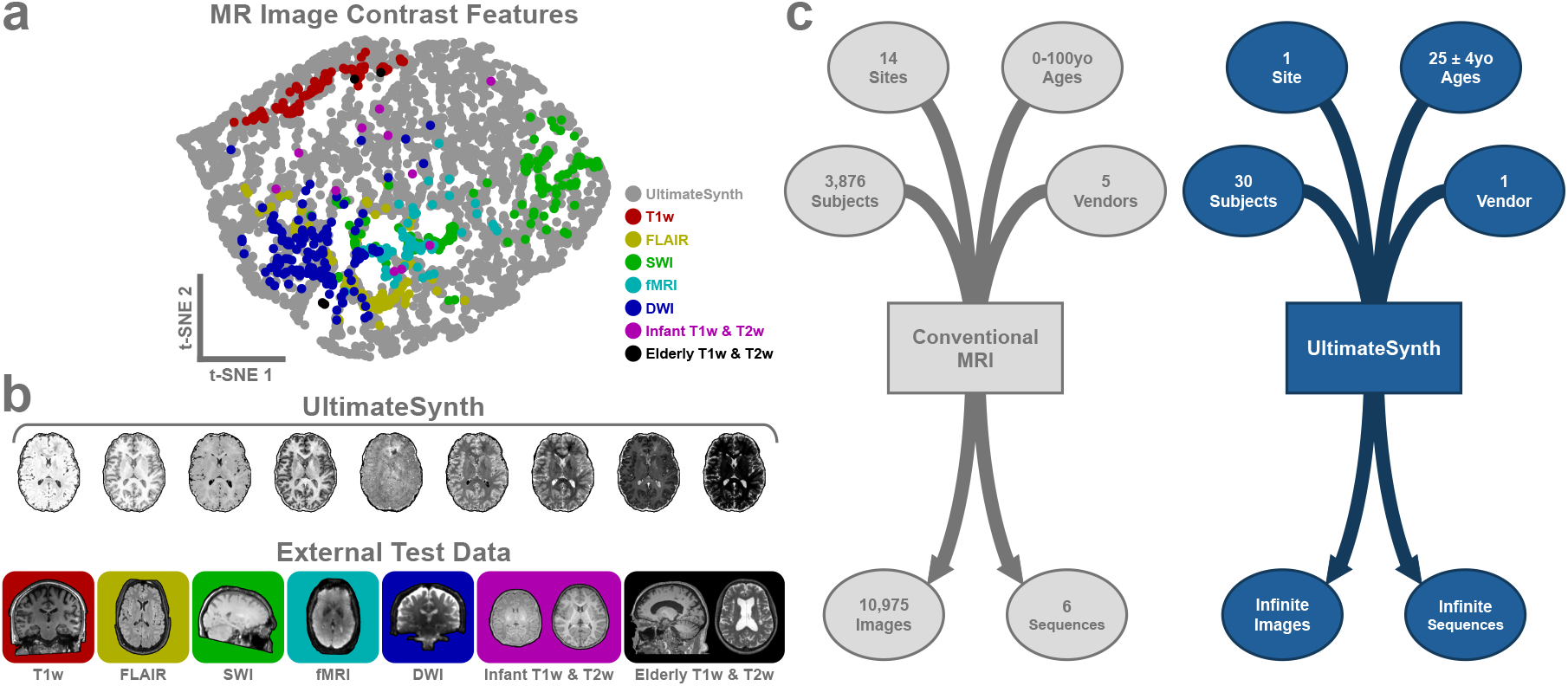
Pan-contrast completeness of UltimateSynth. **a**, t-SNE^20^ visualization of 15 first-order radiomic features of 3,200 UltimateSynth images (gray) and 575 external test images from the ON-Harmony^24^ (red, yellow, green, teal, blue), Baby Connectome Project^25^ (purple), and FreeSurfer Maintenance^26^ (black) datasets. **b**, Representative images of the datasets in **a**, spanning three major age groups and five MR scan types (T1w, T2w/DWI, FLAIR, SWI, and fMRI). **c**, Comparison between the evaluation dataset and the UltimateSynth training dataset for the segmentation models developed in this study in terms of subjects, sites, ages, and scanner vendors.

### Efficient Labeling for Pan-Contrast Learning

Utilizing UltimateSynth for labeling and emulating pan-contrast data, we trained the off-the-shelf nnU-Net^22^ for two segmentation tasks: Whole-brain segmentation (UBN: UltimateBrainNet) with automated labeling and cerebellum segmentation (UCN: UltimateCerebelumNet) with manual labeling. The goal is to show that broad generalizability can be achieved with minimal labeling.

UBN was trained on 30 subjects, each contributing 600 images (100 per one of six major contrasts) at 1 mm isotropic resolution, totaling 18,000 images (see Efficient Contrast Sampling). Figure 2c shows an example of the inter-tissue contrast diversity in these training images. Each image is augmentated via random skull stripping, extended neck simulation, bias field corruption, random reorientation, random flipping, random translation, random resolution downsampling, and random uniform and non-uniform noise (see Training Data Augmentation). Reference labels were generated automatically by applying FreeSurfer—a widely used whole-brain segmentation tool often treated as a silver-standard alternative to manual labeling—to multiple UltimateSynth images of the same anatomy with varied contrasts (as in Figure 2b; results shown in Figure 4a). Across 30 subjects with 8 FreeSurfer segmentations each, the average percent disagreement from majority (PDM) was 4.43 ± 10.69%, and label volume variation (LVV) was 6.17 ± 7.31% for non-background labels (Figures 4b–c). PDM captures voxel-wise disagreement with a majority vote; LVV reflects label-wise volume variability (see Evaluation Metrics). FreeSurfer performance can degrade with certain contrasts, with white matter volumes deviating up to 19.65% from majority. To mitigate contrast-dependent inconsistencies, majority voting was used per subject to create a more robust ground truth for training.

**Fig. 4.**
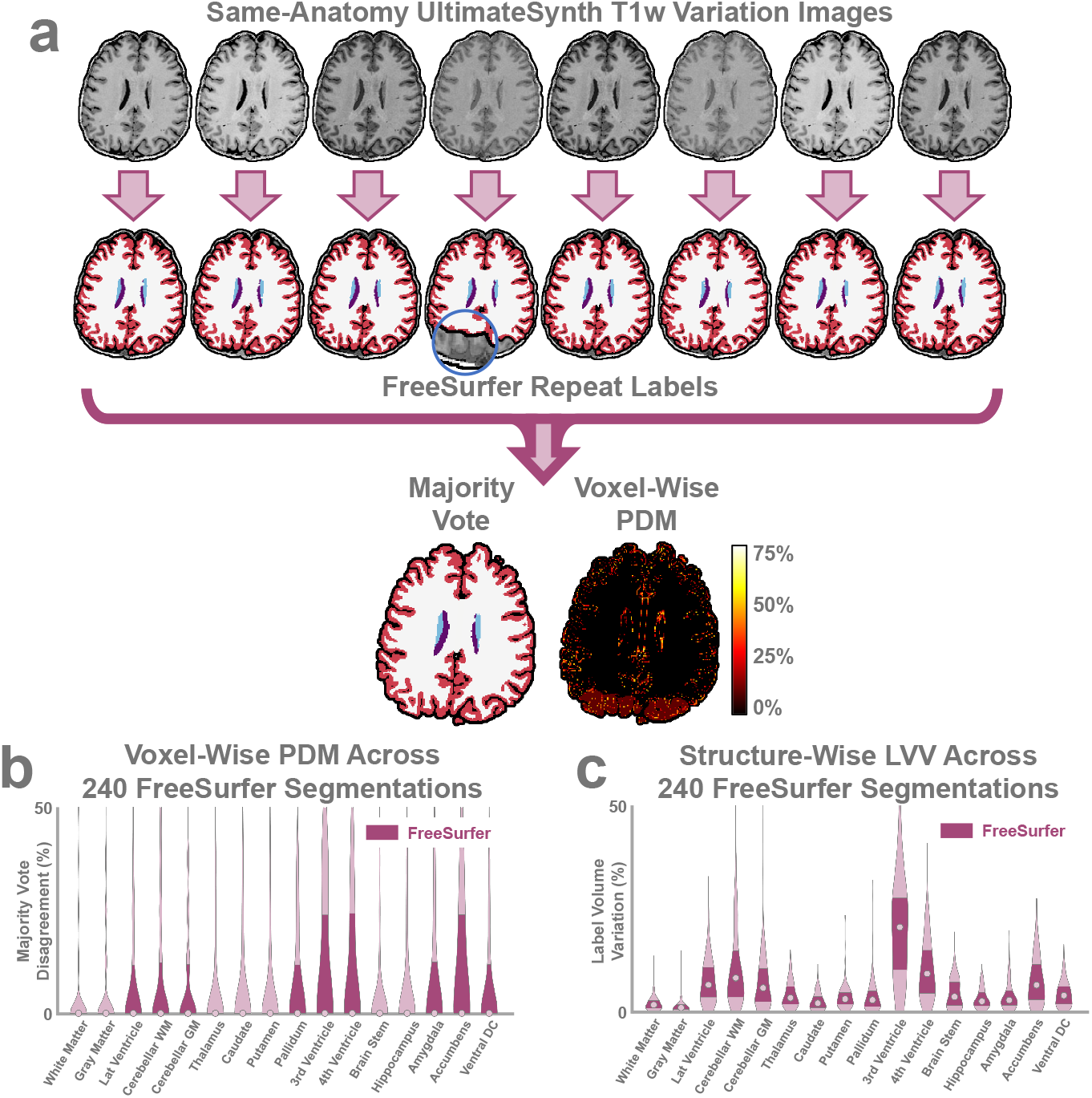
á Concensus labeling using UltimateSynth. **a**, UltimateSynth generates multiple images per subject, enabling combined FreeSurfer segmentations to reduce contrast-related errors and major failures (blue circle). **b**, Voxel-wise percent disagreement from majority (PDM) across 240 segmentations (30 subjects × 8 repeats), indicating labeling consistency (lower is better). **c**, Structure-wise label volume variation (LVV) across the same 240 segmentations.

UCN was trained using 2 subjects manually labeled with UltimateSynth images with cerebellum-enhanced contrasts. Reference labels were created by manually segmenting cerebellar white and gray matter on contrasts customized to highlight the WM-GM boundary (Figure 2b). For each subject, 600 images at 0.5 mm isotropic resolution were generated (100 per one of six major contrasts). The same augmentations used for UBN were applied.

### Pan-Contrast Performance Benchmarking

UltimateSynth provides a platform for comprehensive performance and generalizability assessment of AI networks prior to model dissemination. This platform allows for rigorous assessment of models, regardless of whether they are trained with or without UltimateSynth, across a broad spectrum of image contrasts. The flexible control over image generation provided by UltimateSynth also facilitates investigation of potential failure modes across the full contrast landscape.

For comprehensive pan-contrast evaluation, we generated 150,000 UltimateSynth images from 8 test subjects (18,750 per subject). Movie S1 presents a broad sample of 756 images from a single subject, covering all unique contrast classes (Contrast Classes). Using this dataset, we compared segmentation performance between UBN and the state-of-the-art (SOTA) SynthSeg^15^ (Figure 5a).

**Fig. 5.**
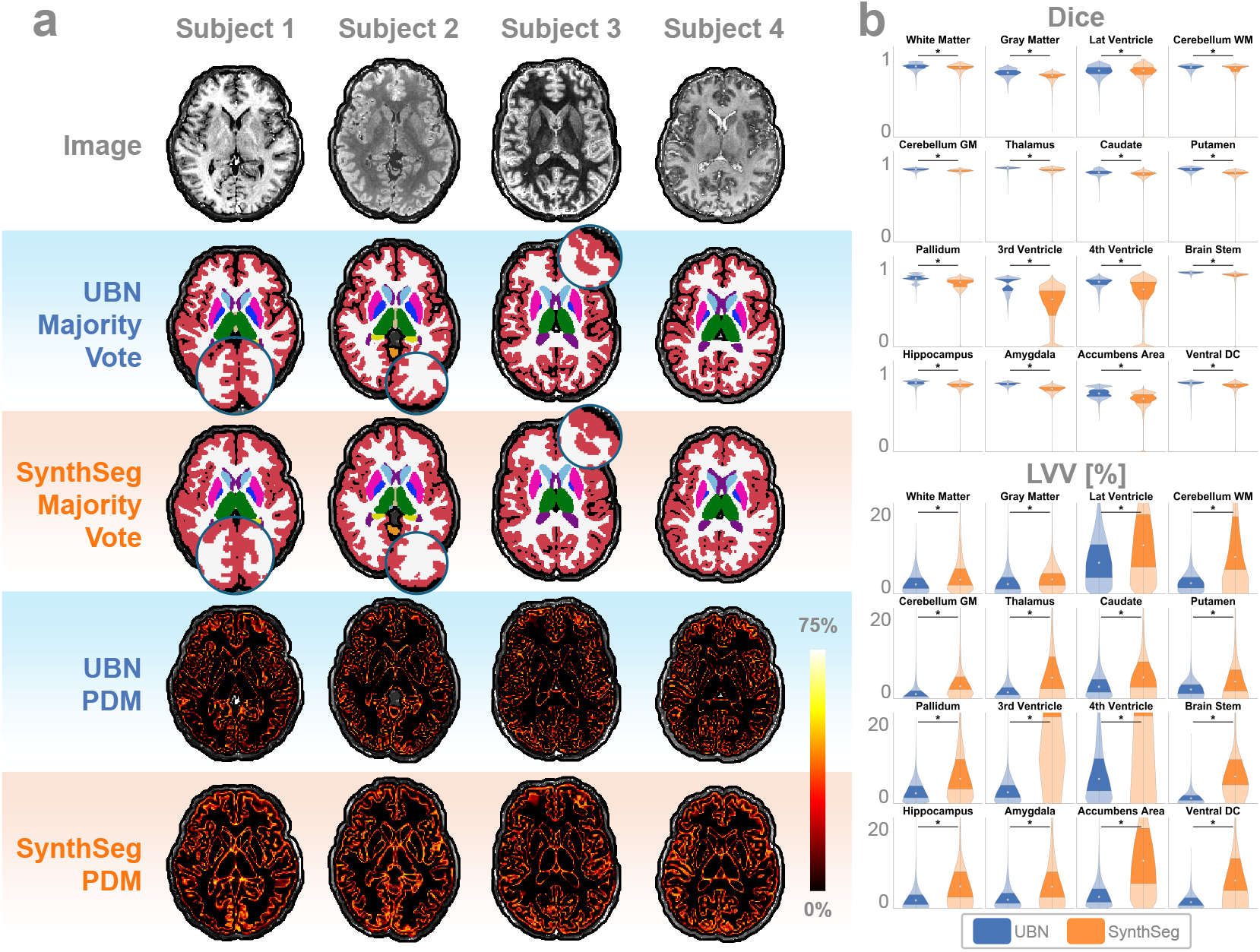
Segmentation performance evaluation using 150,000 unique images generated by UltimateSynth. **a**, Four example subjects used in the evaluation of UBN and SynthSeg. For each subject, 18,750 unique contrasts were generated, spanning a uniformly distributed range of image quality across the UltimateSynth contrast landscape. Voxel-wise PDM maps highlight regions where each method produces inconsistent segmentations across contrast variations. **b**, Structure-specific Dice and LVV scores were computed across all test images. UBN significantly outperforms SynthSeg in Dice accuracy against FreeSurfer labels and maintains under 8% median volume variation across all structures (*P* < .05).

Across all 150,000 segmentations, UBN achieved a subject-average PDM of 7.38 ± 1.35% for non-background labels, outperforming SynthSeg’s 10.78 ± 2.76%. The largest improvements were observed near partial volume boundaries, especially in the basal ganglia. PDM computation was based on method-specific consensus labels generated by voxel-wise majority voting across all segmentations for each subject. Dice scores, computed against FreeSurfer consensus labels, further highlight UBN’s advantage. UBN attained a label-average Dice score of 0.83 ± 0.07, compared to 0.76 ± 0.19 for SynthSeg—a 7-point improvement (Figure 5b; Table S1). UBN yielded higher Dice scores in 87.2% of all label instances (Figure S1). UBN also demonstrated more consistent anatomical accuracy, achieving a lower label-average LVV of 3.35 ± 3.75% versus 16.41 ± 33.28% for SynthSeg, reflecting a 13.06% improvement (Figure 5b; Table S1). These performance gains persisted across a variety of MR scan types (Figure S2b; Table S2). Figure S2a illustrates UBN’s consistent advantage across images with good, medium, and poor contrasts. In its worst case, UBN still achieved a label-average Dice score of 0.59, with errors primarily at ambiguous white matter boundaries, while deep brain structures remained intact. In contrast, SynthSeg’s worst-case Dice score dropped to 0.10, with anatomically implausible segmentations.

### Intra-Subject Consistency Across Sites, Scanners, and Contrasts

To assess UBN’s intra-subject repeatability across scanner and site variations, we used the publicly available ON-Harmony dataset^24^, which comprises 560 3D MRI scans from 10 subjects. The dataset includes data from 6 scanners across 5 sites and 3 vendors. It spans 5 MRI modalities at 4 resolutions: T1weighted MRI and FLAIR (1 mm isotropic), diffusion-weighted imaging (DWI, 2 mm isotropic), susceptibility-weighted imaging (SWI, 0.8 × 0.8 × 3 mm^3^), and resting-state functional MRI (fMRI, 2.4 mm isotropic).

Figure 6 summarizes the segmentation results of UBN, SynthSeg, and PhysSeg on 160 T1weighted and FLAIR images. Despite mild to severe contrast variations across repeat scans of the same subject, UBN consistently outperformed SynthSeg and PhysSeg for all contrast types and scanners (Figure 6a). Quantitative results are shown in Figure 6b and Table S3. These findings indicate that UBN yields *harmonized* measurements that remain consistent across diverse imaging conditions. Dice scores from co-registered T1w and FLAIR scans of the same anatomy show that UBN outperforms SynthSeg by 1.4 points, with mean scores across all labels of 0.864 ± 0.06 versus 0.850 ± 0.06. While PhysSeg achieves a similar mean Dice score of 0.81 ± 0.06, it is limited to a 3-label segmentation scheme and supports only T1w images. UBN also yields significantly lower LVV than both SOTA methods in 13 out of 16 anatomical structures, as shown in Table S4. Compared to SynthSeg, UBN reduces LVV by nearly 4%, achieving 3.42 ± 3.14 versus 7.34 ± 8.23.

**Fig. 6.**
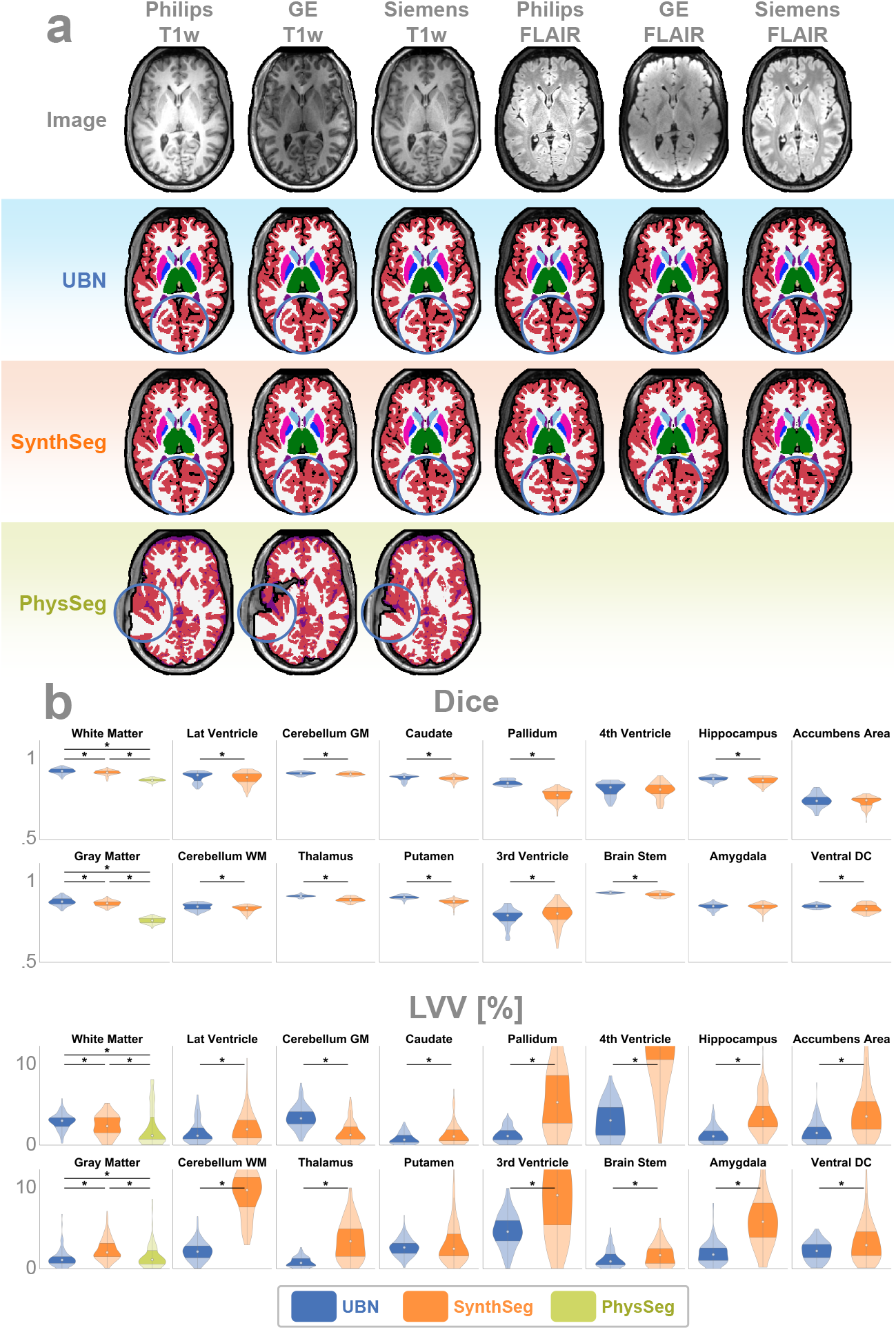
Intra-subject consistency across sites, scanners, and contrasts evaluated using the ON-Harmony dataset. **a**, Comparison of UBN, SynthSeg, and PhysSeg on a representative subject scanned using scanners from three major vendors and with two contrasts. PhysSeg results for FLAIR images are excluded, as the model supports only T1-weighted images. SynthSeg struggles to capture fine cortical folding in the occipital lobe, while PhysSeg shows major errors along the pial surface and in the putamen (blue circles). UBN provides more accurate segmentations across all six contrasts in both regions. **b**, Quantitative evaluation of 160 images spanning ten subjects, three vendors, five sites, six scanners, and two contrasts. UBN demonstrates less than 5% LVV across all 16 labels, with statistically significant LVV improvements over the SOTA for 13 of 16 structures and significant Dice improvements for 12 of 16 structures (*P* < .017 for white matter and gray matter, < .05 for all other labels). PhysSeg is notably limited by its restriction to three-tissue labeling.

We further evaluated UBN on additional MRI modalities, including SWI (Figure S3, Table S5), DWI (Figure S4, Table S6), and fMRI (Figure S5, Table S7). These scans differ widely in resolution and quality, and often present geometric distortions and cropped fields of view (FOVs). Despite not being explicitly trained on such variations, UBN significantly outperforms SynthSeg in maintaining label consistency across repeated scans.

### Lifespan Segmentation Consistency

UBN’s robustness to age-related anatomical variations were investigated using a total of 4,273 T1w images and 4,112 T2w images from subjects ranging from 6 days old to 100 years old. These images were assembled from several datasets: the Baby Connectome Project (BCP)^25^, the Human Connectome Project (HCP)^27, 28^, and the Healthy Brain Network (HBN)^29^, and varied in resolution: 0.7 mm isotropic (HCP), 0.8 mm isotropic (BCP, HCP, and HBN), and 1 mm isotropic (HBN). We observe both high agreement between T1w and T2w volume trajectories (Figure 7a). The trajectories are in line with healthy aging: rapid growth in infancy and adolescence followed by adulthood plateaus and then gradual atrophy in advanced age. Equivalent SynthSeg segmentation trajectories are shown in Figure S6. For the 4,090 image pairs with intra-subject intra-session paired T1w and T2w data, we directly compare T1w-T2w segmentation volume differences in Figure 7b for UBN and SynthSeg, noting that perfect segmentations should yield a 0% volume difference. UBN has a T1w-T2w volume mean absolute error (MAE) of 3.48 ± 3.06% across all labels, a 5.13% improvement over SynthSeg’s 8.61 ± 13.95% MAE. MAE values for specific labels are reported in Table S8.

**Fig. 7.**
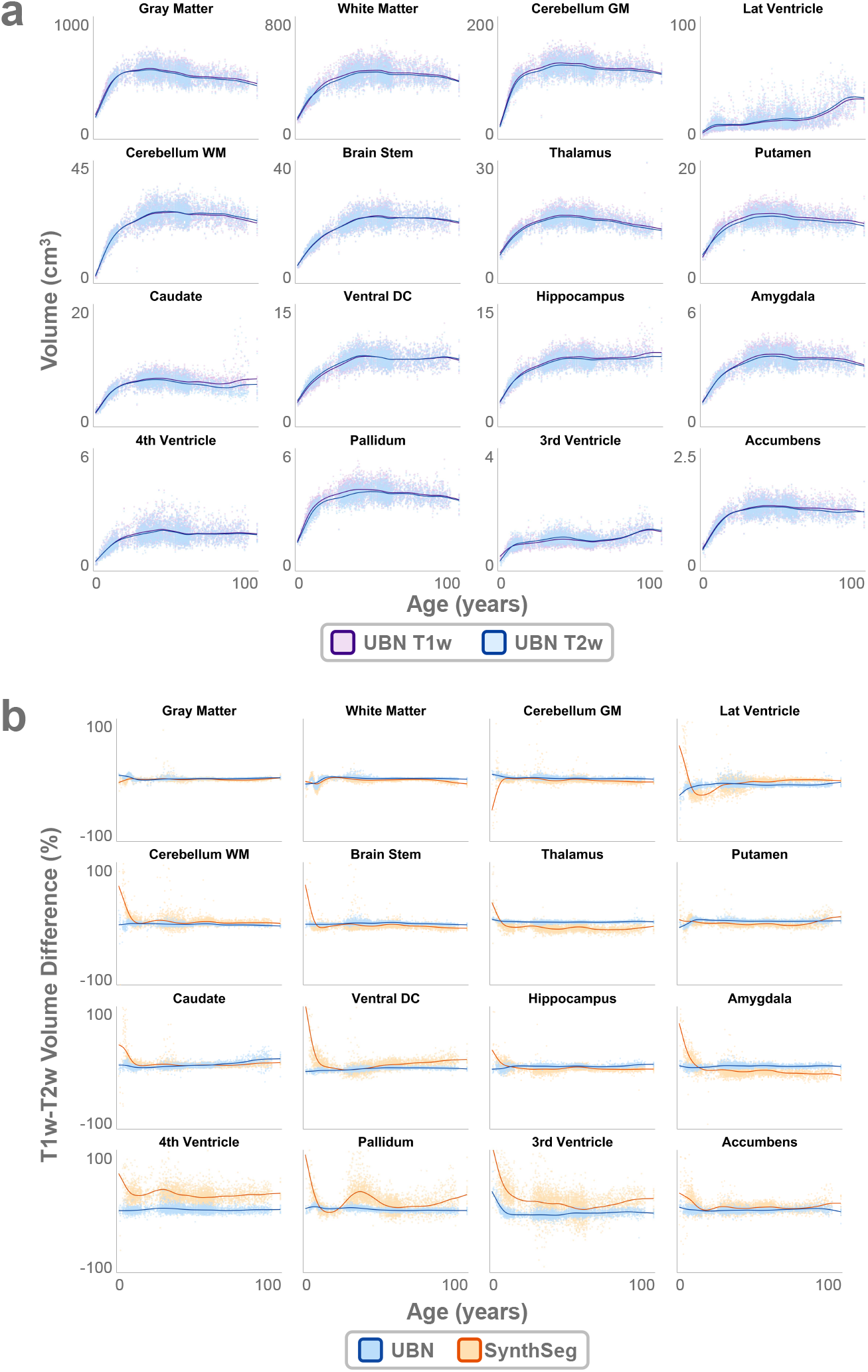
Segmentation across the human lifespan. **a**, UBN label volumes from segmentations of 8,385 T1w and T2w images of subjects ranging from 6 days old to 100 years old. Volume trends, plotted separately for T1w (purple) and T2w (blue) images, indicate high intra-method agreement. **b**, Intra-subject label volume differences between 4,090 paired T1w and T2w images for UBN (blue) and SynthSeg (orange) across the human lifespan. While SynthSeg exhibits significant variation in label volumes in several structures like the cerebellum, thalamus, and pallidum, especially at young ages, UBN has near 0-percent volume difference trends.

Infant brains are especially challenging cases due to unique intra-tissue and cross-age intensity changes caused by gradual myelination processes^30^. We apply UBN, SynthSeg, and Infant FreeSurfer^31^ (infantFS) to a subset of the subjects from the BCP dataset in Figure 8. In contrast to the obvious fail cases for non-UltimateSynth based methods for the 17-day-old image, we note appreciable consistency in the repeated segmentation of anatomical features by UBN, despite age-related contrast changes driven by myelination.

**Fig. 8.**
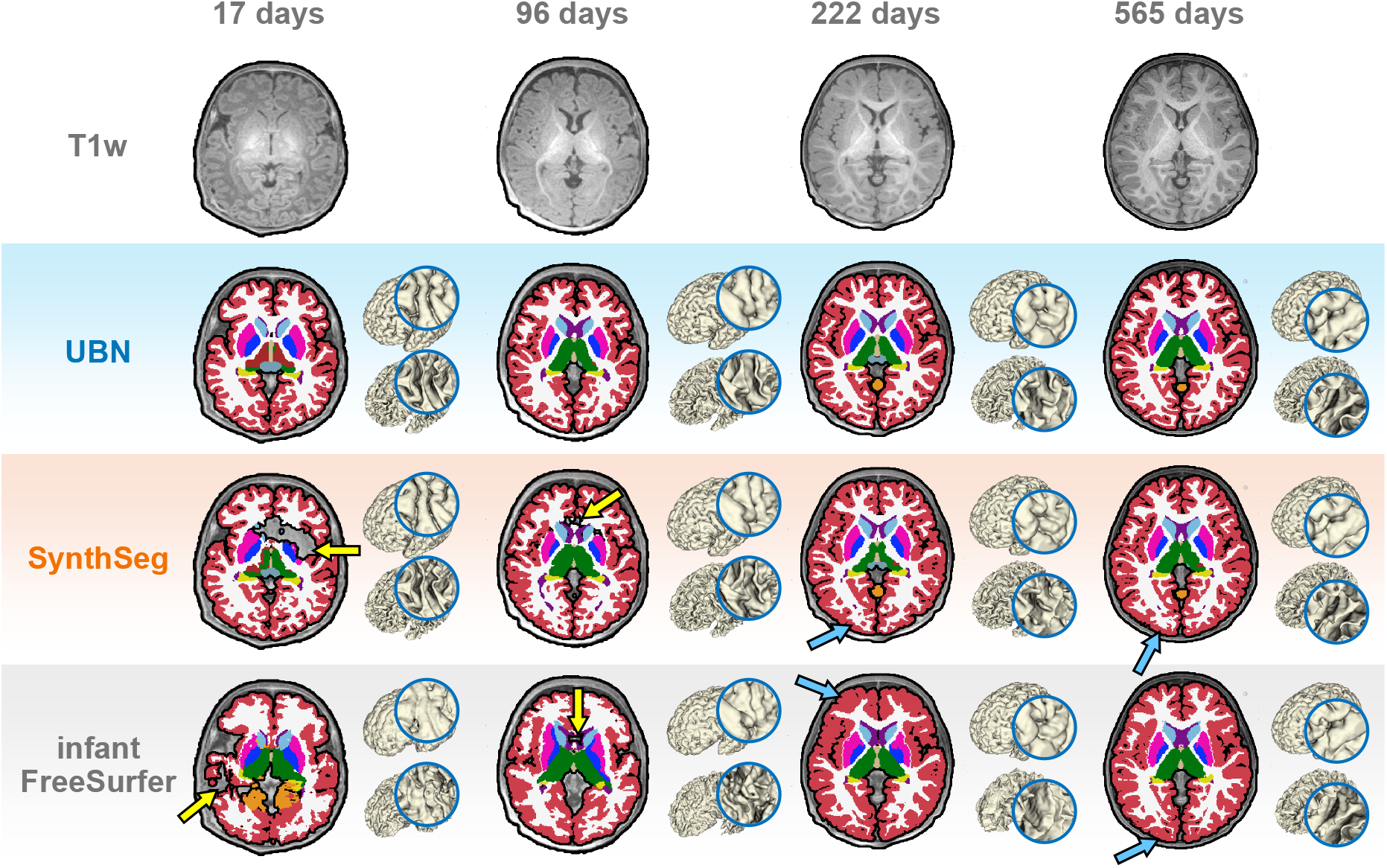
Segmentation generalizability across age-related tissue contrast variations. Longitudinal segmentation consistency from birth to 18 months. UBN consistently segments narrow sulci across time, whereas SynthSeg and Infant FreeSurfer often produce markedly different tissue boundaries in response to age-related contrast changes, despite identical underlying anatomy (blue arrows). These inconsistencies are evident in both the pial (top) and white matter (bottom) surface reconstructions over time. At very early ages, such as 17 and 96 days, existing methods may also produce severe segmentation errors (yellow arrows).

### Generalizability to Low-Field MRI

UBN was evaluated on 2,030 low-resolution MR images from the open-source M4Raw dataset^32^, acquired at 0.3T with a resolution of 0.94 × 1.23 5 mm^3^. The dataset includes 990 T1w, 624 T2w, and 416 FLAIR images from 208 unique subjects. For each subject, LVV was computed across repeat scans and contrasts for both UBN and SynthSeg. As shown in Figure 9a, UBN segmentations better preserve key anatomical structures—such as the pial surface, corpus callosum, and internal capsule—compared to SynthSeg. Quantitatively (Figure 9b, Table S9), UBN achieves a mean LVV improvement of 3.07% over SynthSeg, with statistically significant gains in 15 out of 16 regions.

**Fig. 9.**
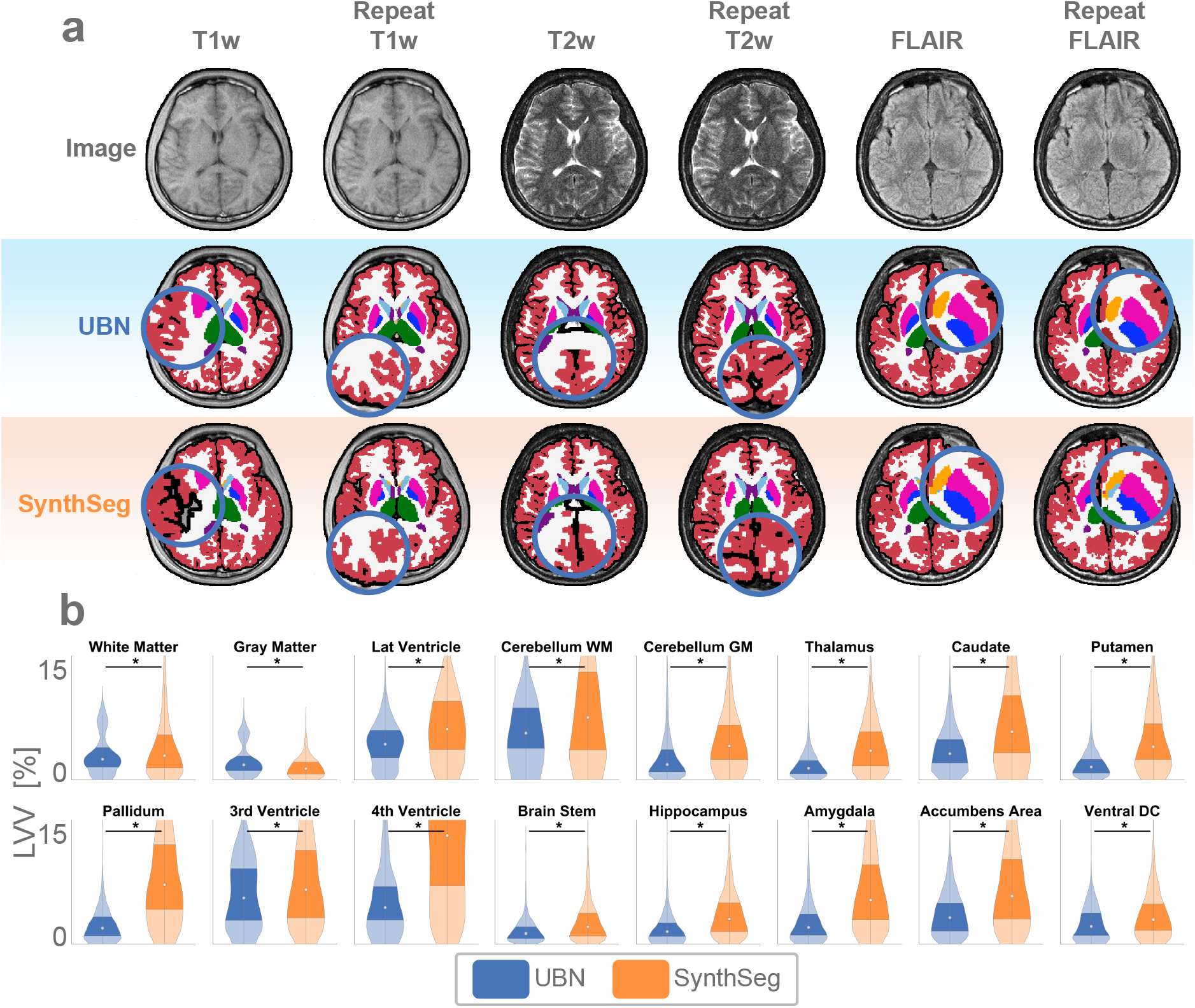
Segmentation of low-field MRI. **a**, A representative subject with repeat 0.3T scans at 0.94 × 1.23×5 mm^3^ resolution for T1w, T2w, and FLAIR sequences, along with corresponding segmentations from UBN and SynthSeg. Compared to SynthSeg—which often exhibits major errors such as pial surface erosion, corpus callosum discontinuity, and loss of the internal capsule white matter (blue circles)—UBN produces visibly more accurate results. **b**, Quantitative evaluation of label volume variation (LVV) across 2,030 T1w, T2w, and FLAIR images from the same subjects. UBN shows significantly lower LVV than SynthSeg in 15 out of 16 labels.

### Lifespan Segmentation of the Cerebellum

Comparing UCN with the state-of-the-art CerebNet^33^ on 8,385 cross-lifespan images, Figure 10a illustrates UCN’s finer segmentation of cerebellar white matter, in contrast to CerebNet’s more conservative estimates. This difference is also reflected in the cross-lifespan volume trajectories shown in Figure 10b, where CerebNet tends to overestimate gray matter and underestimate white matter volumes relative to UCN. Quantitatively, UCN achieves strong consistency between T1w and T2w segmentations, with a mean absolute volume difference of just 2.44 ± 2.28% across the lifespan (Table S10).

**Fig. 10.**
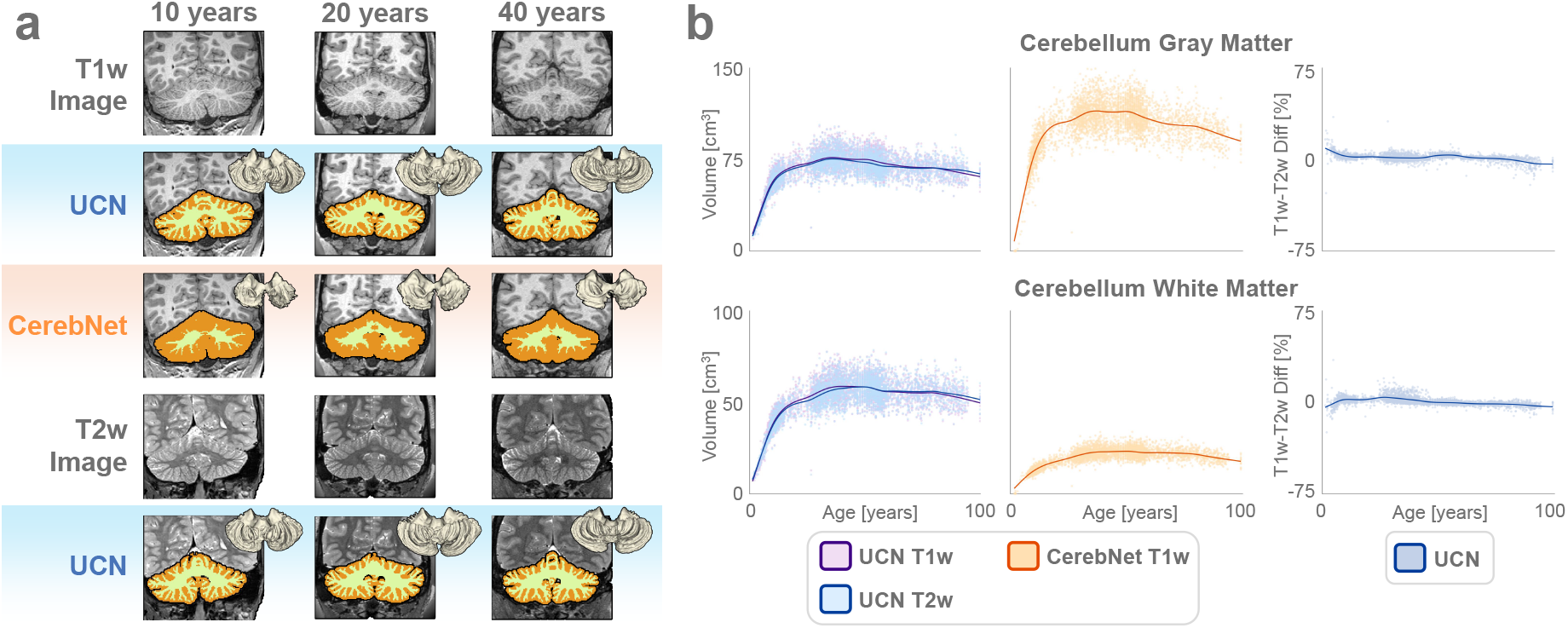
Lifespan cerebellar segmentation. **a**, Qualitative comparison of UCN and CerebNet in the axial view, showing reconstructed cerebellar white matter surfaces. **b**, Label volume trajectories across the human lifespan for UCN and CerebNet. A total of 4,273 T1w and 4,112 T2w images were segmented using UCN, while CerebNet was applied only to the T1w images due to its modality limitation. CerebNet consistently underestimates white matter and overestimates gray matter volumes relative to UCN. UCN also shows strong cross-modality consistency, with near-zero percent volume differences between paired T1w and T2w segmentations.

### Ablation Study

Two experiments were conducted to evaluate how different components of UBN’s training contribute to its ability to generalize across image contrasts: (i) the number of anatomical training examples, and (ii) the extent of data augmentation.

To evaluate how the number of unique anatomical examples affects model performance, we trained six neural networks using the same set of 6,400 UltimateSynth-generated image contrasts. These images were derived from varying numbers of unique subjects—ranging from 1 to 32—resulting in models denoted UBN 1 through UBN 32. All models were tested on 160 T1w and FLAIR images described in Intra-Subject Consistency Across Sites, Scanners, and Contrasts. Models trained with fewer unique anatomies (e.g., UBN 1) struggled with unexpected anatomical variations during testing, particularly with enlarged lateral ventricles (Figure S7a, yellow arrows). Performance improved as the number of training subjects increased, with a mean Dice gain of 0.05 from UBN 1 to UBN 32. However, improvements plateaued beyond 16 subjects, with only a 0.003 difference in mean Dice between UBN 16 and UBN 32 (Figure S7b, Table S11). t-SNE^20^ visualization (Figure S8b) showed that pan-contrast generalization was similar for UBN 1 and UBN 32, indicating that while anatomical diversity enhances robustness to structural variability, it has little impact on contrast generalization. This aligns with findings in Lifespan Segmentation Consistency, where UltimateSynth produced consistent segmentation performance across diverse subjects of all ages.

To evaluate the impact of data augmentation on model performance, we re-trained UBN with substantially reduced augmentation. In this ablated version, key components were removed from the image synthesis pipeline, including random skull stripping, extended neck simulation, image translation, rotation, flipping, resolution downsampling, and synthetic background noise. Both the original and ablated models were tested on the 160 T1w and FLAIR images from IntraSubject Consistency Across Sites, Scanners, and Contrasts. As shown in Figure S9a, the ablated model produces clear misclassifications of brain labels in non-brain regions. This degradation is also reflected quantitatively in Figure S9b and Table S12, where the ablated model performs significantly worse than UBN on key structures such as white matter, lateral ventricles, and the caudate nucleus. However, for labels confined to brain regions—like the ventral diencephalon and hippocampus—both models perform similarly. These results suggest that while data augmentation (e.g., bias field and background noise simulation) helps prevent errors outside the brain, UltimateSynth’s simulation is sufficiently robust to support accurate segmentation within the brain, even with minimal augmentation.

## DISCUSSION

We introduced UltimateSynth—a versatile framework that integrates MR physics and tissue physiology to generate a wide spectrum of realistic MR contrasts without repeated scanning. UltimateSynth supports the labeling, emulation, and deployment stages of AI development. We demonstrated that UltimateSynth facilitates the generation of labeled data for training networks that are highly generalizable, capable of covering 150,000 unique contrasts with exceptionally low volumetric variability under 4%.

Our approach fundamentally differs from existing simulation methods by directly modeling spin dynamics based on intrinsic tissue properties and real MR physics—without relying on predefined pulse sequences. This removes constraints imposed by scanner hardware, conventional imaging protocols, and sequence-specific approximations, enabling comprehensive yet realistic coverage of the MR contrast landscape. Unlike traditional simulators that require a separate model for each pulse sequence, our unified framework can generate a broad range of image contrasts—including T1-weighted, T2-weighted, FLAIR, SWI, and beyond. This generalizability supports both accurate emulation of clinical protocols and flexible contrast optimization, whether globally across the image or locally within specific regions. As a result, we move beyond simple protocol mimicry or random image generation to provide a powerful platform for contrast design, training data synthesis, efficient labeling, and robust benchmarking across diverse applications.

We enable efficient pan-contrast image synthesis through a novel decomposition of MRI signal dictionaries into a contrast parameter subspace that is independent of anatomy. In this continuous contrast landscape, each point represents a distinct image contrast. Because the subspace is derived using orthogonal SVD components, movement in any direction reliably results in a noticeable change in image contrast. This property is distinct from explicit sequence modeling, where exponential signal decay can lead to visually similar images—even when parameters vary widely at extreme values. The completeness of this contrast space is demonstrated in Figure S8a: 3,200 randomly generated UltimateSynth images fully capture the range of first-order radiomic contrast intensity features found in 570 test images across five pulse sequence types and multiple age groups.

The primary motivation for simulating the full pan-contrast MR landscape with UltimateSynth is to address a key challenge in clinical AI deployment: test-time out-of-distribution failure. MRI contrasts vary widely due to differences in hardware, pulse sequences, and visualization, and while most cases may appear typical, model robustness is most critically tested under rare or edge-case scenarios. By training across a deliberately all-inclusive contrast spectrum— including extreme and atypical cases—we equip models to handle real-world variability. Including extreme contrasts is therefore essential for building resilient, generalizable, and safe models across diverse scanners, populations, and clinical scenarios.

### Customizable Synthesizer for Efficient Labeling

UltimateSynth tackles two key challenges in medical image labeling: label scarcity and contrast-induced uncertainty. By enabling automatic propagation of a single label set across infinite unique contrasts, it eliminates the need for repeated labeling, boosts data diversity, and ensures consistency across contrasts to enhance model robustness. UltimateSynth adapts image contrast to match standard labeling tools (e.g., T1w for FreeSurfer), enabling automated, scalable label generation. It also improves anatomical visibility with tailored contrasts, simplifying manual labeling and facilitating the creation of new labels. Finally, Label-time contrast augmentation allows multiple labeling outcomes for the same structure, enabling reliability assessment and refinement through techniques like majority voting. This improves label accuracy, boosts model performance, and supports the creation of higher-quality ground truth for model learning and asssessment.

### Platform for Deployment Benchmarking

UltimateSynth enables comprehensive benchmarking of AI models by providing diverse, customizable imaging contrasts—addressing the lack of a standardized platform for evaluating generalizability across contrasts. Traditional assessments require large, labeled datasets spanning varied contrasts, which are impractical due to manual annotation demands. Misalignments from patient positioning or acquisition variability further distort evaluations, especially for small structures. UltimateSynth overcomes these limitations with a unified platform that uses qMR maps to generate perfectly aligned, multi-contrast test images, enabling accurate, structure-consistent evaluation. We validated this by segmenting over 18,000 contrast images per test subject (see Pan-Contrast Performance Benchmarking), allowing detailed analyses of volumetric and voxel-level segmentation errors to identify model limitations. UltimateSynth also supports performance analysis as a function of image quality, as shown in Figures S1 and S2a.

### Pan-Contrast Generalization with Limited Training Data

UBN and UCN demonstrate strong generalizability with minimal training data, highlighting UltimateSynth’s effectiveness in building robust AI models. UBN was trained on just 30 healthy volunteers (mean age 25.6 ± 4.3 years) yet showed consistent performance across diverse conditions and the full human lifespan (see Supplementary Figures S10–S14). UBN also generalized well to independent datasets with substantial variability—spanning three MR vendors, six scanners, five sites, five scan types, and varying resolutions and field strengths—demonstrating strong robustness without the need for extensive retraining or tuning.

### Simulation Time and Computational Constraints

During training, both UBN and UCN used 600 images per subject—100 for each contrast type defined in Contrast Classes. Generating these images takes about one hour per subject: approximately 24 seconds to load the data and map it to the contrast landscape, and 1.5–6 seconds to generate an image depending on augmentation. These times do not account for parallelization, which is feasible across images or subjects. Each subject requires around 3.5 GB of RAM, mainly for the SVD-based contrast decomposition. Although UltimateSynth is fast enough to support on-the-fly synthesis, our offline uniform sampling of the contrast landscape already provides sufficient variability to enable pan-contrast generalization, as demonstrated by experimental results.

Training the UBN model involved a large number of images and the use of five-fold crossvalidation, presenting a computational bottleneck. However, by parallelizing across five NVIDIA V100 GPUs—one per fold—training was completed in under six days. No further tuning was needed, thanks to the model’s robust pan-contrast generalizability. The most time-consuming step was repeatedly running the FreeSurfer pipeline (about 9 hours per run) per subject to generate consensus labels. Fortunately, FreeSurfer processing is fully parallelizable across subjects and contrasts, so the total time for all 240 segmentations can be significantly reduced depending on available computational resources.

#### Limitations

While UBN and UCN are compatible with any MR image, UltimateSynth currently depends on the availability of qMR tissue property maps. Although such maps are not yet common in public datasets, Magnetic Resonance Fingerprinting (MRF) is an FDA approved method for simultaneously acquiring qMR maps, and thus the widespread data availability of qMR maps is expected to increase substantially in the near future. As shown in Ablation Study, even a small number of subjects can suffice to train useful models with UltimateSynth. Moreover, recent advances in estimating qMR maps from standard MRI may soon enable UltimateSynth to work with any existing scans.

#### Future Work

UltimateSynth’s current use in healthy brain MRI illustrates just one example of its broader potential. Future applications include pathological imaging, other anatomical regions, and even different species. By generating high-quality, diverse, and well-labeled datasets across a wide range of contrasts, UltimateSynth supports the development of versatile models that can serve as foundations for advanced tasks such as super-resolution, spatial normalization, and anomaly detection. This enables the creation of contrast-agnostic, robust AI tools that bridge research and clinical practice, advancing diagnostics, treatment planning, and personalized medicine.

## METHODS

### Tissue Property Maps

MR Fingerprinting (MRF)^34^, an efficient and clinically feasible technique^34–36^, was used to obtain the tissue quantitative maps from 40 healthy volunteers (25.6 ± 4.3 years of age) at a single site^37^. Scanning was approved by the local Institutional Review Board and were conducted using a 3T Siemens Prisma scanner with a 20-channel head coil. Prior to scanning, all participants provided written informed consent. The tissue property maps from each subject included co-registered quantitative T1, T2, and proton density maps, all simultaneously acquired from a single MRF scan. Three-dimensional head scans were acquired with whole brain coverage using an axial acquisition, a 300 × 300 × 144 mm^3^ field of view, 1 mm isotropic image resolution, and a scan time of 10 minutes. A fast B1 mapping scan (20 seconds) was acquired from the same subject to measure the transmit field inhomogeneity. Quantitative T1, T2, and proton density maps were generated using a low-rank algorithm. 32 of these subjects were allocated for training UBN and UCN via UltimateSynth image simulation. Two subjects were excluded from UBN’s training set due to excessive motion artifacts that degraded the quality of their qMR maps, resulting in a final training dataset of 30 subjects. The remaining 8 subjects were reserved for test-time analysis(see Pan-Contrast Performance Benchmarking).

### Traveling Human Data

The ON-Harmony^24^ traveling heads dataset was utilized for repeatability testing purposes. The data consists of ten subjects (33.2 ± 9.9 years of age) each with between 6 and 11 scan sessions across three vendors, five sites, and six scanners, for a total of 560 combined T1-weighted, FLAIR, SWI, DWI, and fMRI images. The data were downloaded from the OpenNeuro public database (https://openneuro.org/datasets/ds004712). Intra-subject anatomical scans were rigidly co-registered for subsequent evaluation using FSL’s FLIRT^38–40^. Original resolutions were 1 mm isotropic for T1-weighted and FLAIR, 2 mm isotropic for DWI, 4. mm isotropic for fMRI, and 0.8 × 0.8 × 3 mm^3^ for SWI. All images were resampled to 1 mm isotropic resolution using linear interpolation as a preprocessing step.

### Lifespan Data

A total of 8,385 T1w and T2w images of subjects ranging from 6 days old to 100 years old were acquired from five public datasets to investigate UBN and UCN performance across the human lifespan: the Baby Connectome Project (BCP)^25^, Healthy Brain Network (HBN)^29^, Human Connectome Project Development (HCPD)^27^, Human Connectome Project Young Adult (HCPYA)^28^, and the Human Connectome Project Aging (HCPA)^27^. Subject demographics and image acquisition parameters for each dataset are summarized below:

- **BCP**: 969 T1w images and 819 T2w images were used. The mean age at time of scan was 1.52 ± 1.19 years. 206 subjects were female, 192 male, and 1 unknown. Images were acquired with a 3T Siemens Prisma using a 32 channel head coil. T1w images were acquired sagittaly with 208 slices, TR = 2,400 ms, TE = 2.24 ms, flip angle = 8^°^, matrix size of 320 × 320, and 0.8 mm isotropic resolution. T2w images were acquired sagittaly with 208 slices, TR = 3200 ms, TE = 564 ms, variable flip angle, matrix size of 320 × 320, and 0.8 mm isotropic resolution.
- **HBN**: 771 T1w and 760 T2w images were used. The mean age at time of scan was 11.73 ± 3.53 years. 275 subjects were female, 459 male, and 37 unknown. Images were acquired with 1.5T Siemens Avanto and 3T Siemens Trio scanners. T1w images were acquired with either 176 or 224 slices, TR = 2730/2500 ms, TE = 1.64/3.15/2.9/2.88 ms, flip angle = 7/8^°^, matrix size of 256 × 256 or 320 × 320, and 1 mm isotropic or 0.8 mm isotropic resolution. T2w images were acquired with either 176 or 224 slices, TR = 3200 ms, TE = 564/565 ms, flip angle = 120^°^, matrix size of 256 × 256 or 320 × 320, and 1 mm isotropic or 0.8 mm isotropic resolution.
- **HCPD**: 651 T1w and 651 T2w images were used. The mean age at time of scan was 14.44 ± 4.06 years. 351 subjects were female and 301 were male. Images were acquired with a 3T Siemens Prisma using a 32 channel head coil. T1w images were acquired with 208 slices, TR = 2500 ms, TE = 1.8/3.6/5.4/7.2 ms, flip angle = 8^°^, matrix size of 320 × 300, and 0.8 mm isotropic resolution. T2w images were acquired with 208 slices, TR = 3200 ms, TE = 564 ms, matrix size of 320×300, and 0.8 mm isotropic resolution.
- **HCPYA**: 1158 T1w and 1158 T2w images were used. The mean age at time of scan was 28.80±3.70 years. Images were acquired with a 3T ConnectomScanner. T1w images were acquired with 260 slices, TR = 2400 ms, TE = 2.14 ms, flip angle = 8^°^, matrix size of 260×311, and 0.7 mm isotropic resolution. T2w images were acquired with 260 slices, TR = 3200 ms, TE = 564 ms, variable flip angle, matrix size of 260×311, and 0.7 mm isotropic resolution.
- **HCPA**: 724 T1w and 724 T2w images were used. The mean age at time of scan was 60.37 ± 15.73 years. 405 subjects were female and 319 were male. Images were acquired with a 3T Siemens Prisma using a 32 channel head coil. T1w images were acquired with 208 slices, TR = 2500 ms, TE = 1.8/3.6/5.4/7.2 ms, flip angle = 8^°^, matrix size of 320 × 300, and 0.8 mm isotropic resolution. T2w images were acquired with 208 slices, TR = 3200 ms, TE = 564 ms, matrix size of 320×300, and 0.8 mm isotropic resolution.

### Low-Field Data

The M4Raw^32^ low-field low-resolution dataset was used for testing. The dataset consists of 208 subjects and a total of 2,030 images: 990 T1w, 624 T2w, and 416 FLAIR. All images were acquired at 0.3T and 0.94 × 1.23×5 mm^3^ resolution. T1w spin echo images were acquired with a matrix size of 256 × 195, TR = 500 ms, TE = 18.4 ms, field of view of 240×240 mm^2^, and 18 slices with slice thickness of 5 mm. T2w fast spin echo images were acquired with a matrix size of 256×195, TR = 5500 ms, TE = 128 ms, field of view of 240 × 240 mm^2^, and 18 slices with slice thickness of 5 mm. FLAIR spin echo images were acquired with a matrix size of 256 × 198, TR = 7500 ms, TE = 98 ms, TI = 1655 ms, field of view of 240 × 240 mm^2^, and 18 slices with slice thickness of 5 mm. The data were downloaded from the public M4Raw data repository (https://zenodo.org/records/8056074). All images were linearly interpolated to 0.8 mm isotropic resolution before inference.

### Elderly Data

T1-weighted and T2-weighted MRI scans for two elderly subjects (76 years of age and 72 years of age) demonstrating appreciable lateral ventricle enlargement were obtained from the open-source FreeSurfer Maintenance Dataset^26^ for visualization of contrast features. The data were downloaded from the OpenNeuro public database (https://openneuro.org/datasets/ds004958) and intra-subject scans were rigidly co-registered using FLIRT^38–40^.

### Physics-Informed Contrast Generation

In nuclear magnetic resonance (NMR) and magnetic resonance imaging (MRI), the acquired signal is determined by both intrinsic tissue properties (e.g., relaxation times, proton density, diffusion, and hemodynamics) and extrinsic imaging factors (e.g., static magnetic field (B0), radiofrequency (RF) field (B1), and acquisition timings (echo time, TE; repetition time, TR)). Here, we applied the classic Bloch equations^41^ to synthesize diverse image contrasts, covering the full range of combinations of imaging factors (initial magnetization, RF pulses, TR, TE, and phase) from a set of tissue properties (T1, T2, and proton density). Specifically, the spin magnetization *M*_*r*_∈ ℝ ^3^ at a given echo time (TE) is described by the state equation^42^ of a single isochromat *r*:

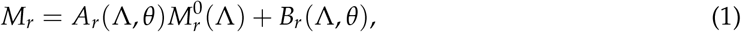

where 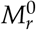 is the initial magnetization prior to the RF excitation. This magnetization is in the form of [0 0 *M*_*z*_]^T^, where the magnetization along the z-axis, *M*_*z*_ ∈ [− 1, 1], can be modified by various preparation mechanisms, such as T1 inversion, T2 preparation, or diffusion preparation pulses. We simulated the 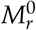 magnetization from an inversion preparation pulse with randomly varied inversion time (TI) without physical limitation. *θ* represents a set of tissue properties (e.g., T1, T2, and proton density) and experiment-specific parameters (e.g., B1 inhomogeneity), whereas Λ comprises imaging factors, including RF pulses and TE times. *A*_*r*_ is a system matrix *A*_*r*_(Λ, *θ*) = *R*(T1, T2, TE)*Q*(*α, ϕ*), with *R*(T1, T2, *t*) modeling spin relaxation:

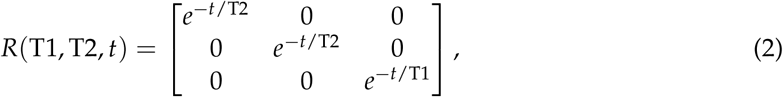

and *Q*(*α, ϕ*) modeling RF excitation:

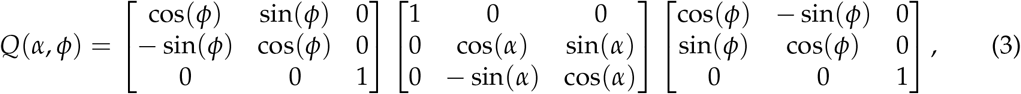

where *α* is the RF excitation flip angle and *ϕ* is the RF pulse phase. Lastly, *B*_*r*_(Λ, *θ*) is the input matrix:

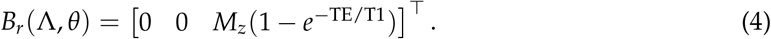

In practice, we simulate *θ*_1_, *θ*_2_, …, *θ*_*j*_ as a series of unique combinations of T1 and T2, where T1 ∈ (0 5000] ms and T2 ∈ (0 2000] ms, and we simulate Λ_1_, Λ_2_, …, Λ_*k*_ as a series of unique combinations of *α*, TI, and TE, where *α* ∈ (0 360] degrees, TI ∈ [1 5000] ms, and TE [1 400] ms. This allows for the construction of a two-dimensional voxel intensity dictionary matrix *D*^*⋆*^, where rows encode all combinations of *θ* values and columns encode all combinations of Λ imaging factors:

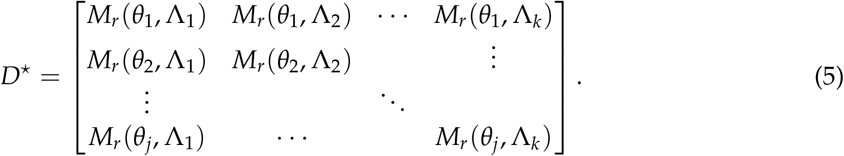

Since network training was based on the signal magnitude, the signal phase was not encoded in the dictionary. However, MR is sensitive to parts per million (p.p.m.) level deviations in the B0 field, and so the signal phase induced from field inhomogeneities can be simulated in the dictionary in the future as well. Ranges were chosen for the imaging factors to ensure that magnetization at TE for any tissue type spans the full range from -1 to 1. A total of 122,080,000 dictionary entries were generated in 1.1 seconds on a standard desktop computer. The relative GM-WM-CSF contrasts for UltimateSynth dictionary entries with unique TI-TE combinations and a fixed 90 degree RF pulse are visualized as a 2D plot in Figure S15.

### Efficient Contrast Sampling

To efficiently sample the contrast landscape, we applied singular value decomposition (SVD) with truncation^43^ to the mean-subtracted matrix *D*^*⋆*^ to extract the principal MR contrasts:

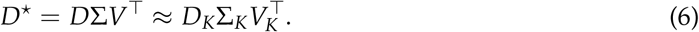

Here, *K* = 10 singular values were retained, preserving over 99.95% of the total energy. The truncated matrix *D*_*K*_ encodes the top *K* eigencontrasts and defines a reduced coordinate system, with rows representing T1 and T2 combinations from 0 to 5,000 ms. A new MR contrast *λ* can be synthesized by specifying a coefficient vector *c*_*λ*_ ∈ ℝ^*K*^, and computing

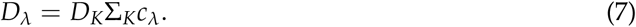

Voxel intensities for a subject’s synthetic image are then retrieved from *D*_*λ*_ using voxel-wise T1-T2 index look-up. By randomly varying *c*_*λ*_, an unlimited number of realistic MR contrasts can be generated without rerunning simulations. To ensure physical plausibility, the elements of *c*_*λ*_ are empirically constrained to the range [−1, 1].

We sample each dimension of the contrast landscape uniformly, allocating more samples to dimensions with larger singular values as given by Σ_*K*_. In the *i*-th dimension, we generate *r*_*i*_ linearly spaced points in the range [−1, 1], where

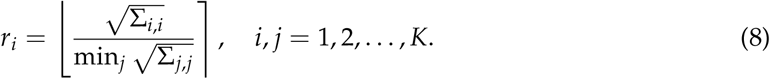

and rounds a number to the nearest integer. A coefficient vector *c*_*λ*_ is then formed by combining one sample from each of the *K* dimensions.

To ensure contrast diversity during training, we sample across the full range of contrast quality within each of the six contrast classes (see Contrast Classes). Specifically, to sample *H* contrasts from a given class, we first compute the distribution of multi-tissue contrast scores for that class. We then divide this distribution into *H* percentile-based bins and draw one sample from each bin.

### Contrast Metrics

To measure relative contrast among tissues ℒ = {WM, GM, CSF}, we first define representative tissue properties: WM (T1 = 850 ms, T2 = 40 ms), GM (T1 = 1,400 ms, T2 = 70 ms), and CSF (T1 = 3,000 ms, T2 = 300 ms). Alternatively, subject-specific values can be estimated by computing the median T1 and T2 within each tissue class from a segmentation map. Given voxel intensities *I*_L_ for tissue classes ℒ under a specific set of imaging parameters, we compute the inter-tissue intensity distances as

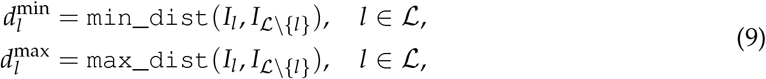

where min_dist and max_dist represent the minimum and maximum absolute intensity differences between a given tissue and all other tissues in the set. Defining the overall contrast range as _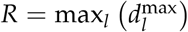_, we introduce two contrast metrics:

- **Single-tissue contrast score:** Measures how well a single tissue type *l* is distinguished from the others:

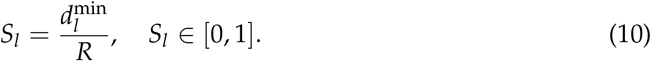

- **Multi-tissue contrast score:** Measures overall separability across all tissue types:

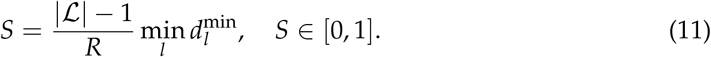

### Contrast Classes

Six distinct contrast classes can be defined based on relative intensities: (1) *I*_WM_ *> I*_GM_ *> I*_CSF_, (2) *I*_WM_ *> I*_CSF_ *> I*_GM_, (3) *I*_GM_ *> I*_WM_ *> I*_CSF_, (4) *I*_GM_ *> I*_CSF_ *> I*_WM_, (5) *I*_CSF_ *> I*_GM_ *> I*_WM_, and (6) *I*_CSF_ *> I*_WM_ *> I*_GM_. Based on these orderings, the 150,000 images used in Figure 5 and Figure S2 were named as follows:

- **T1-weighted:** 25,000 images with the ordering *I*_WM_ *> I*_GM_ *> I*_CSF_,
- **T2-weighted:** 25,000 images with the ordering *I*_CSF_ *> I*_GM_ *> I*_WM_,
- **FLAIR:** 25,000 images with the ordering *I*_GM_ *> I*_WM_ *> I*_CSF_,
- **Others:** 75,000 images spanning the remaining three orderings.

### Data Processing and Consensus Labeling

The T1-weighted region of the eigencontrast matrix *D* (WM > GM > CSF) was uniformly sampled, and the multi-tissue contrast score *S* (Eq. 11) was computed at each point. This region was split at the 60th percentile of *S* to separate higher and lower contrast regions. Eight coefficient vectors from the higher-contrast subset were randomly selected to generate eight T1-weighted images per subject, each with varying inter-tissue contrast levels. Each synthetic T1-weighted image was intensity-normalized between its 1st and 99th percentiles and processed with FreeSurfer’s recon-all pipeline^21^. Parcellation outputs were condensed into 16 regions of interest. For each subject, voxel-wise majority voting across the eight segmentations produced a consensus label map, enhancing labeling consistency and reliability.

### Labeling Outside the Brain

To reduce confusion during network training, the 16 condensed FreeSurfer labels were augmented with two additional anatomical labels: a 17th label for unlabeled nonzero-intensity voxels within the brain mask (subarachnoid space), and an 18th for nonzero voxels outside the brain mask (skull, scalp, and other tissues). Two additional labels captured minor and major background noise. At test time, no assumptions were made about skull presence in the input image.

### Label Post-Processing

At test time, UBN and UCN segmentations were refined using connected-component analysis to suppress extraneous labels outside the brain. For UBN, labels 1–17 were binarized, and the largest connected component was used to mask the final segmentation. For UCN, the same was done using only labels 1 and 2 (cerebellar white and gray matter).

### UBN Training

For each training subject, 600 UltimateSynth images at 1 mm isotropic resolution were generated by uniformly sampling the contrast space *D* as detailed in Efficient Contrast Sampling, using FreeSurfer reference labels as described in Data Processing and Consensus Labeling. This yielded a total training set of 18,000 images covering a wide range of MR contrasts. A 3d_fullres nnUNetv2 model was trained using a combined cross entropy and Dice loss against eight-time majority-voted FreeSurfer labels. Training ran for 600 epochs on the CWRU HPC using an Nvidia V100 GPU (32GB memory), requiring 58 hours in total.

### UCN Training

For two training subjects with manual reference labels, 600 UltimateSynth images each were generated by uniformly sampling the contrast space *D* as detailed in Efficient Contrast Sampling. These images were then linearly upsampled to 0.5 mm isotropic resolution, resulting in 1,200 training images. A 3d_fullres nnUNetv2 was trained for 1,000 epochs on the CWRU HPC using an Nvidia V100 GPU (32GB memory). The model optimized a combined cross entropy and Dice loss between predictions and manual annotations, completing training in 69.4 hours.

### Training Data Augmentation

To improve network performance, training data augmentation steps was randomly applied to generated UltimateSynth images, including (i) skull stripping and neck synthesis, (ii) bias field corruption, 3) reorientation, (iii) flipping, (iv) translation, (v) resolution downsampling, (vi) uniform background noise, and (vii) non-uniform background noise. In total, synthesizing an image took ∼ 1.5 seconds with minimal augmentation and ∼ 6 seconds with maximal augmentation.

- **Skull Stripping and Neck Synthesis**: For one-third of randomly selected images, the original skull was left unchanged. For another one-third, skull stripping was applied using masks derived from FreeSurfer reference labels, retaining only brain structures and cerebrospinal fluid (CSF)/meninges. In the remaining one-third of UltimateSynth training images, extended neck and spine regions were synthetically simulated. The MRF maps used for model training typically have a limited field of view, excluding most of the throat, shoulders, and neck. This omission can lead to inference issues when those regions are present in test-time data. To address this, we first fitted each training subject with a deformable head and neck atlas^44^ to obtain anatomical labels for non-brain structures, including spinal cord, blood vessels, air, muscles, mucosa, skin, and bone beyond the original imaging field. During neck simulation, we computed the mean and standard deviation of each atlas label within the field of view of the initial image. These statistics were then propagated beyond the original field of view using voxel-wise Gaussian sampling within each atlas-defined region, generating a simplified synthetic neck and upper spine.
- **Bias Field**: Intensity inhomogeneity or shading results from scanner-related variations such as magnetic field inhomogeneities, differences in radiofrequency coil sensitivity, and other imaging imperfections. The associated bias field is both scanner- and subject-dependent, making it difficult to model and correct accurately. To simulate realistic bias fields for U-Net training, we adopted the method described in SynthSeg^15^. Specifically, we generated a low-resolution 3 × 3 × 3 bias volume *B*^′^ with voxel-wise intensities sampled from a zeromean Gaussian distribution, 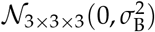, where the standard deviation *Σ*_B_ for each voxel was drawn from a uniform distribution 𝒰 (0, *b*_B_). To obtain the final smooth bias field *B*, we upsampled *B*^′^ using cubic spline interpolation and applied a voxel-wise exponential transform. This yielded multiplicative intensity scaling values constrained to the range [0, 1]. The synthetic bias field *B* was applied to the clean image *G* to generate a bias-corrupted version: *G*_*B*_ = *G* × *B*.
- **Reorientation**: Each image is randomly rotated in 3D around the three spatial axes—[0, 0, 1], [0, 1, 0], and [1, 0, 0]. For each axis, the rotation angle is independently selected from {0^°^, 90^°^, 180^°^, 270^°^} with equal probability.
- **Flipping**: For each of the three spatial axes, there is a 50% chance that the image is randomly flipped along that axis.
- **Translation**: Along each spatial axis, a random translation is applied by sampling from a uniform distribution scaled to the image dimension. The minimum and maximum translation values are constrained to ensure that the ground truth brain labels of interest remain within the image field of view.
- **Downsampling**: In 50% of training images, linear downsampling is applied along one randomly selected spatial axis, with equal probability of reducing the resolution to either 2 mm or 4 mm in that direction. Among these downsampled images, 50% are further downsampled along a second randomly chosen axis, and of those, 50% are downsampled along the third axis as well. Each downsampling step independently chooses between 2 mm and 4 mm resolution. This procedure results in (i) 50% of images remaining at 1 mm isotropic resolution, (ii) 25% downsampled along one axis, (iii) 12.5% downsampled along two axes, and (iv) 12.5% downsampled along all three axes, producing resolutions as low as 4 mm isotropic. Following the simulation of noise in the next two steps, all images are resampled back to 1 mm isotropic resolution to match the resolution of the reference labels.
- **Uniform Noise**: In 50% of training images, random Gaussian background noise is added. To determine the noise level, we first compute the 5th percentile of voxel intensities within the reference labels of interest in the synthetic image. A Gaussian distribution is then defined with both mean and standard deviation equal to this 5th percentile value, and a single sample from this distribution is used to set the noise level for the image. For each voxel outside the reference labels, background noise is added by drawing a value from a zero-mean Gaussian distribution with the sampled standard deviation and summing it with the original voxel intensity.
- **Non-Uniform Noise**: In 50% of images, non-uniform background noise is added. This is done by selecting random regions defined by randomly generated rectangular prisms that exclude foreground voxels (as defined by the reference labels of interest). Each selected region is filled with Gaussian noise, where the mean and standard deviation are randomly drawn from values between the 5th and 95th percentiles of voxel intensities within the reference labels. After adding a noise region, there is a 50% chance of repeating the process and adding an additional noise block, and a 50% chance of stopping. This introduces variable, spatially localized noise patterns across the background.

### Evaluation Metrics

Segmentation performance was assessed using the volume-weighted SørensenDice coefficient:

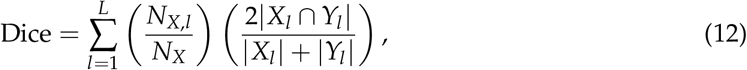

where *L* is the number of labels, *N*_*X*_ is the total number of labeled voxels in *X*, and *N*_*X,l*_ is the count of voxels labeled *l* in *X. X*_*l*_ and *Y*_*l*_ denote the binary masks for label *l* in *X* and *Y*.

Voxel-wise inconsistency was measured using Percent Disagreement from the Majority Vote (PDM). For a set of *J* images with *N* voxels labeled from {1, …, *L*}, the majority label at voxel *n* is and the corresponding disagreement is

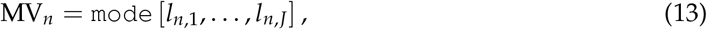

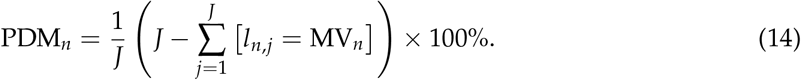

Label-wise variability was quantified using Label Volume Variation (LVV). The volume of label *l* in image *j* is

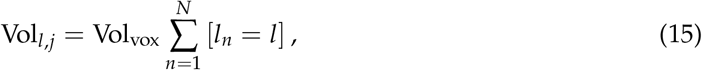

where Vol_vox_ is the voxel volume. The mean volume across *J* images is

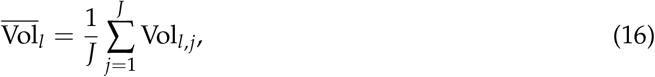

and the LVV is

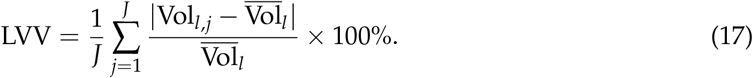

### Visualization of Pan-Contrast Completeness

We analyzed voxel intensity features from the 3,200 whole-brain training images used for UBN, along with 575 test images. For each image, non-background voxels were normalized between the 1st and 99th percentiles, and intensities were linearly scaled to the [0, 255] range to enable fair comparison across images. Using either the reference or UBN segmentation (if reference was unavailable), we computed 15 first-order intensity-based radiomics features for each of the 16 anatomical labels, as defined by PyRadiomics^45^. These include statistics such as energy, percentiles, mean, standard deviation, skewness, and kurtosis—yielding 240 features per image. Applying t-SNE to these features reveals patterns in inter-tissue intensity profiles, effectively characterizing image contrast across the dataset (Supplementary Figure S8).

### Visualization of Quantitative Metrics

Figures 4, 5, 6, 9, S2, S3, S4, S5, S7, and S9 present quantitative segmentation performance using violin plots^46, 47^ to visualize data distributions. In each plot, the median is shown as a circle, while the top and bottom of the dark-shaded region indicate the first and third quartiles. The width of the violin reflects data density along the vertical axis—wider sections represent more frequent values, and narrower sections indicate sparser data.

## Statistical Analysis

For all intra-subject repeat segmentations, paired Wilcoxon signed-rank tests were used to compare Dice and LVV scores between UltimateSynth-based models and SOTA alternatives. Statistical significance was defined as *P* < .05 and Bonferroni correction was applied for comparisons involving more than two methods. Performance results are reported as Mean ± Standard Deviation.

## Supplementary Information

The manuscript contains supplementary material.

## Acknowledgements

This work was supported in part by the United States National Institutes of Health (NIH) under grants R01 CA269604 (D. Ma), R01 CA282516 (D. Ma), R01 NS109439 (D. Ma), R01 EB035160 (P.-T. Yap), R01 MH125479 (P.-T. Yap), R01 EB008374 (P.-T. Yap), R01 NS134849 (P.-T. Yap), and R01 HD112923 (D. Ma and P.-T. Yap). The authors would like to thank Juan Eugenio Iglesias for his comments on the initial version of the manuscript.

## Author Contributions

R.A.: methodology, software, investigation, visualization, writing — original draft, writing — review and editing. K.M.H.: resources, data curation, visualization, investigation, writing — review and editing. W.Z.: methodology, resources. S.H.: resources, data curation. W.L.: resources, data curation. S.A.: resources, data curation, visualization. D.M.: conceptualization, methodology, supervision, funding acquisition, investigation, validation, writing — review and editing. P.-T.Y.: conceptualization, methodology, supervision, funding acquisition, investigation, validation, writing — review and editing.

## Code Availability

Software used to analyze data and prepare figures for this manuscript includes MATLAB R2024a and the following associated toolboxes: Deep Learning Toolbox 14.5, Image Processing Toolbox 11.6, Parallel Computing Toolbox 7.7, Signal Processing Toolbox 9.1, and the Statistics and machine Learning Toolbox 12.4. Additional software used includes FreeSurfer 7.4.1, FSL 6.0.7.7, nnUNet v2, Violins for MATLAB 0.1, SynthSeg 2.0, PhysSeg 0.1, infantFreeSurfer dev-4a14499, FastSurfer 2.4.2 (CerebNet), Python 3.11.8 and the following associated packages: numpy 1.26.4, sklearn 1.4.1.post1, matplotlib 3.8.3, and h5py 3.10.0.

## Correspondence

Correspondence and requests for materials should be addressed to D.M. and P.-T.Y.

## References

1. Huppertz, H.-J., Kröll-Seger, J., Klöppel, S., Ganz, R. E. & Kassubek, J. Intra- and interscanner variability of automated voxel-based volumetry based on a 3D probabilistic atlas of human cerebral structures. NeuroImage 49, 2216–2224 (2010). URL https://linkinghub.elsevier.com/retrieve/pii/S1053811909011379.

2. Fortin, J.-P., Sweeney, E. M., Muschelli, J., Crainiceanu, C. M. & Shinohara, R. T. Removing inter-subject technical variability in magnetic resonance imaging studies. NeuroImage 132, 198–212 (2016). URL https://linkinghub.elsevier.com/retrieve/pii/S1053811916001452.

3. Hedges, E. P. et al. Reliability of structural MRI measurements: The effects of scan session, head tilt, inter-scan interval, acquisition sequence, FreeSurfer version and processing stream. NeuroImage 246, 118751 (2022). URL https://linkinghub.elsevier.com/retrieve/pii/S1053811921010235.

4. Filippi, M. et al. Interscanner variation in brain MRI lesion load measurements in MS: Implications for clinical trials. Neurology 49, 371–377 (1997). URL https://www.neurology.org/doi/10.1212/WNL.49.2.371.

5. Clark, K. A., Woods, R. P., Rottenberg, D. A., Toga, A. W. & Mazziotta, J. C. Impact of acquisition protocols and processing streams on tissue segmentation of T1 weighted MR images. NeuroImage 29, 185–202 (2006).

6. Carré, A. et al. Standardization of brain MR images across machines and protocols: bridging the gap for MRI-based radiomics. Scientific Reports 10, 12340 (2020). URL https://www.nature.com/articles/s41598-020-69298-z.

7. Vasiliuk, A., Frolova, D., Belyaev, M. & Shirokikh, B. Limitations of out-of-distribution detection in 3D medical image segmentation. Journal of Imaging 9, 191 (2023). URL https://www.mdpi.com/2313-433X/9/9/191.

8. Mårtensson, G. et al. The reliability of a deep learning model in clinical out-of-distribution MRI data: A multicohort study. Medical Image Analysis 66, 101714 (2020). URL https://linkinghub.elsevier.com/retrieve/pii/S1361841520300785.

9. Benoit-Cattin, H., Collewet, G., Belaroussi, B., Saint-Jalmes, H. & Odet, C. The SIMRI project: a versatile and interactive MRI simulator. Journal of Magnetic Resonance 173, 97–115 (2005). URL https://linkinghub.elsevier.com/retrieve/pii/S1090780704003908.

10. Liu, F., Velikina, J. V., Block, W. F., Kijowski, R. & Samsonov, A. A. Fast realistic MRI simulations based on generalized multi-pool exchange tissue model. IEEE Transactions on Medical Imaging 36, 527–537 (2017). URL http://ieeexplore.ieee.org/document/7676360/.

11. Tanenbaum, L. et al. Synthetic MRI for clinical neuroimaging: Results of the magnetic resonance image compilation (MAGiC) prospective, multicenter, multireader trial. American Journal of Neuroradiology 38, 1103–1110 (2017).

12. Borges, P. et al. Acquisition-invariant brain MRI segmentation with informative uncertainties. Medical Image Analysis 92, 103058 (2024). URL https://linkinghub.elsevier.com/retrieve/pii/S1361841523003183.

13. Jog, A., Hoopes, A., Greve, D. N., Van Leemput, K. & Fischl, B. PSACNN: Pulse sequence adaptive fast whole brain segmentation. NeuroImage 199, 553–569 (2019). URL https://linkinghub.elsevier.com/retrieve/pii/S1053811919304252.

14. Tobin, J. et al. Domain randomization for transferring deep neural networks from simulation to the real world. In IEEE/RSJ International Conference on Intelligent Robots and Systems, 23–30 (2017).

15. Billot, B. et al. SynthSeg: Segmentation of brain MRI scans of any contrast and resolution without retraining. Medical Image Analysis 86, 102789 (2023). URL https://linkinghub.elsevier.com/retrieve/pii/S1361841523000506.

16. Billot, B. et al. Robust machine learning segmentation for large-scale analysis of heterogeneous clinical brain MRI datasets. Proceedings of the National Academy of Sciences 120, e2216399120 (2023). URL https://pnas.org/doi/10.1073/pnas.2216399120.

17. Wells, W., Viola, P., Atsumi, H., Nakajima, S. & Kikinis, R. Multi-modal volume registration by maximization of mutual information. Medical Image Analysis 1, 35–51 (1996).

18. Fischl, B. et al. Whole brain segmentation: Automated labeling of neuroanatomical structures in the human brain. Neuron 33, 41–55 (2002).

19. Kazerouni, A. et al. Diffusion models in medical imaging: A comprehensive survey. Medical Image Analysis 88, 1–22 (2023). URL https://doi.org/10.1016/j.media.2023.102846.

20. Van der Maaten, L. & Hinton, G. Visualizing data using t-SNE. Journal of machine learning research 9 (2008).

21. Fischl, B. FreeSurfer. NeuroImage 62, 774–781 (2012). URL https://linkinghub.elsevier.com/retrieve/pii/S1053811912000389.

22. Isensee, F., Jaeger, P. F., Kohl, S. A. A., Petersen, J. & Maier-Hein, K. H. nnU-Net: a self-configuring method for deep learning-based biomedical image segmentation. Nature Methods 18, 203–211 (2021). URL https://www.nature.com/articles/s41592-020-01008-z.

23. Yang, M. et al. Low rank approximation methods for MR fingerprinting with large scale dictionaries. Magnetic Resonance in Medicine 79, 2392–2400 (2018). URL https://onlinelibrary.wiley.com/doi/10.1002/mrm.26867.

24. Warrington, S. et al. A resource for development and comparison of multimodal brain 3T MRI harmonisation approaches. Imaging Neuroscience 1, 1–27 (2023). URL https://direct.mit.edu/imag/article/doi/10.1162/imag_a_00042/118214/A-resource-for-development-and-comparison-of.

25. Howell, B. R. et al. The UNC/UMN Baby Connectome Project (BCP): An overview of the study design and protocol development. NeuroImage 185, 891–905 (2019). URL https://linkinghub.elsevier.com/retrieve/pii/S1053811918302593.

26. Greve, D. N. & Fischl, B. The FreeSurfer maintenance dataset (2024). URL https://openneuro.org/datasets/ds004958/versions/1.0.0.

27. Harms, M. P. et al. Extending the Human Connectome Project across ages: Imaging protocols for the lifespan development and aging projects. NeuroImage 183, 972–984 (2018). URL https://linkinghub.elsevier.com/retrieve/pii/S1053811918318652.

28. Van Essen, D. C. et al. The WU-Minn Human Connectome Project: An overview. NeuroImage 80, 62–79 (2013). URL https://linkinghub.elsevier.com/retrieve/pii/S1053811913005351.

29. Alexander, L. M. et al. An open resource for transdiagnostic research in pediatric mental health and learning disorders. Scientific Data 4, 170181 (2017). URL https://www.nature.com/articles/sdata2017181.

30. Ahmad, S. et al. Multifaceted atlases of the human brain in its infancy. Nature Methods 20, 55–64 (2023). URL https://www.nature.com/articles/s41592-022-01703-z.

31. Zöllei, L., Iglesias, J. E., Ou, Y., Grant, P. E. & Fischl, B. Infant FreeSurfer: An automated segmentation and surface extraction pipeline for T1-weighted neuroimaging data of infants 0–2 years. NeuroImage 218, 116946 (2020). URL https://linkinghub.elsevier.com/retrieve/pii/S1053811920304328.

32. Lyu, M. et al. M4Raw: A multi-contrast, multi-repetition, multi-channel MRI k-space dataset for low-field MRI research. Scientific Data 10, 264 (2023). URL https://www.nature.com/articles/s41597-023-02181-4.

33. Faber, J. et al. CerebNet: A fast and reliable deep-learning pipeline for detailed cerebellum sub-segmentation. NeuroImage 264, 119703 (2022). URL https://linkinghub.elsevier.com/retrieve/pii/S1053811922008242. xPublisher: Elsevier BV.

34. Ma, D. et al. Magnetic resonance fingerprinting. Nature 495, 187–192 (2013). URL https://www.nature.com/articles/nature11971.

35. Ma, D. et al. Fast 3D magnetic resonance fingerprinting for a whole-brain coverage. Magnetic Resonance in Medicine 79, 2190–2197 (2018). URL https://onlinelibrary.wiley.com/doi/10.1002/mrm.26886.

36. Cao, X. et al. DTI-MR fingerprinting for rapid high-resolution whole-brain T_1_, T_2_, proton density, ADC, and fractional anisotropy mapping. Magnetic Resonance in Medicine 91, 987–1001 (2024). URL https://onlinelibrary.wiley.com/doi/10.1002/mrm.29916.

37. Ma, D. et al. Development of high-resolution 3D MR fingerprinting for detection and characterization of epileptic lesions. Journal of Magnetic Resonance Imaging 49, 1333–1346 (2019). URL https://onlinelibrary.wiley.com/doi/full/10.1002/jmri.26319.

38. Jenkinson, M., Bannister, P., Brady, M. & Smith, S. Improved optimization for the robust and accurate linear registration and motion correction of brain images. NeuroImage 17, 825–841 (2002). URL https://linkinghub.elsevier.com/retrieve/pii/S1053811902911328.

39. Jenkinson, M. & Smith, S. A global optimisation method for robust affine registration of brain images. Medical Image Analysis 5, 143–156 (2001). URL https://linkinghub.elsevier.com/retrieve/pii/S1361841501000366.

40. Greve, D. N. & Fischl, B. Accurate and robust brain image alignment using boundary-based registration. NeuroImage 48, 63–72 (2009). URL https://linkinghub.elsevier.com/retrieve/pii/S1053811909006752.

41. Bloch, F. Nuclear Induction. Physical Review 70, 460–474 (1946). URL https://link.aps.org/doi/10.1103/PhysRev.70.460.

42. Zhao, B. et al. Optimal experiment design for magnetic resonance fingerprinting: Cramér-Rao bound meets spin dynamics. IEEE Transactions on Medical Imaging 38, 844–861 (2019). URL https://ieeexplore.ieee.org/document/8481484/.

43. McGivney, D. F. et al. SVD compression for magnetic resonance fingerprinting in the time domain. IEEE Transactions on Medical Imaging 33, 2311–2322 (2014). URL https://ieeexplore.ieee.org/document/6851901.

44. Puonti, O. et al. Accurate and robust whole-head segmentation from magnetic resonance images for individualized head modeling. NeuroImage 219, 117044 (2020). URL https://linkinghub.elsevier.com/retrieve/pii/S1053811920305309.

45. Van Griethuysen, J. J. et al. Computational radiomics system to decode the radiographic phenotype. Cancer Research 77, e104–e107 (2017). URL https://aacrjournals.org/cancerres/article/77/21/e104/662617/Computational-Radiomics-System-to-Decode-the.

46. Hintze, J. L. & Nelson, R. D. Violin plots: A box plot-density trace synergism. The American Statistician 52, 181–184 (1998). URL http://www.tandfonline.com/doi/abs/10.1080/00031305.1998.10480559.

47. Bechtold, B., Fletcher, P., Seamusholden & Gorur-Shandilya, S. Violin plots for Matlab (2021). URL https://zenodo.org/record/4559847.

