## Supplementary Information for "UltimateSynth: MRI Physics for Pan-Contrast AI"

**Movie S1** `UltimateSynth_Example_Images.gif` displays 756 UltimateSynth images generated for a single subject in the Pan-Contrast Performance Benchmarking experiment. Each image's contrast class, as defined in Contrast Classes, is indicated in the bottom left corner.

**Tab. S1** | Structure-specific Dice and LVV scores across 150,000 UltimateSynth images. UBN outperforms SynthSeg for both metrics and all labels ($P < .001$).

| Structure Label | Mean Volume [%] | UBN Dice | SynthSeg Dice | Dice *P* Value | UBN LVV [%] | SynthSeg LVV [%] | LVV *P*-Value |
|---|---|---|---|---|---|---|---|
| White Matter | 35.24 | **.86 ± .04** | .84 ± .08 | **< .001** | **2.83 ± 2.21** | 5.36 ± 7.97 | **< .001** |
| Gray Matter | 45.69 | **.79 ± .05** | .74 ± .06 | **< .001** | **2.96 ± 2.50** | 4.22 ± 4.11 | **< .001** |
| Lateral Ventricle | 0.73 | **.81 ± .07** | .79 ± .14 | **< .001** | **8.67 ± 6.25** | 16.65 ± 17.87 | **< .001** |
| Cerebellar White Matter | 2.21 | **.85 ± .03** | .77 ± .21 | **< .001** | **2.95 ± 2.11** | 21.81 ± 33.24 | **< .001** |
| Cerebellar Grey Matter | 9.15 | **.89 ± .02** | .84 ± .12 | **< .001** | **1.56 ± 1.66** | 8.21 ± 15.86 | **< .001** |
| Thalamus | 1.34 | **.90 ± .03** | .84 ± .18 | **< .001** | **1.99 ± 1.69** | 14.25 ± 24.81 | **< .001** |
| Caudate | 0.68 | **.85 ± .04** | .81 ± .12 | **< .001** | **3.51 ± 2.81** | 9.94 ± 15.98 | **< .001** |
| Putamen | 1.02 | **.89 ± .03** | .83 ± .10 | **< .001** | **2.54 ± 1.83** | 8.19 ± 13.56 | **< .001** |
| Pallidum | 0.28 | **.84 ± .05** | .76 ± .13 | **< .001** | **3.02 ± 2.34** | 11.31 ± 16.92 | **< .001** |
| 3rd Ventricle | 0.04 | **.75 ± .10** | .51 ± .25 | **< .001** | **3.25 ± 2.50** | 46.82 ± 58.20 | **< .001** |
| 4th Ventricle | 0.11 | **.79 ± .06** | .62 ± .27 | **< .001** | **8.22 ± 7.41** | 41.84 ± 79.62 | **< .001** |
| Brain Stem | 1.68 | **.91 ± .01** | .81 ± .22 | **< .001** | **1.55 ± 1.29** | 18.07 ± 31.25 | **< .001** |
| Hippocampus | 0.74 | **.84 ± .03** | .79 ± .13 | **< .001** | **2.48 ± 2.08** | 11.18 ± 18.00 | **< .001** |
| Amygdala | 0.32 | **.83 ± .02** | .76 ± .08 | **< .001** | **2.73 ± 2.32** | 8.49 ± 11.78 | **< .001** |
| Accumbens Area | 0.12 | **.72 ± .06** | .62 ± .14 | **< .001** | **3.49 ± 2.89** | 19.36 ± 23.30 | **< .001** |
| Ventral Diencephalon | 0.65 | **.83 ± .06** | .76 ± .19 | **< .001** | **1.86 ± 1.52** | 16.80 ± 27.07 | **< .001** |

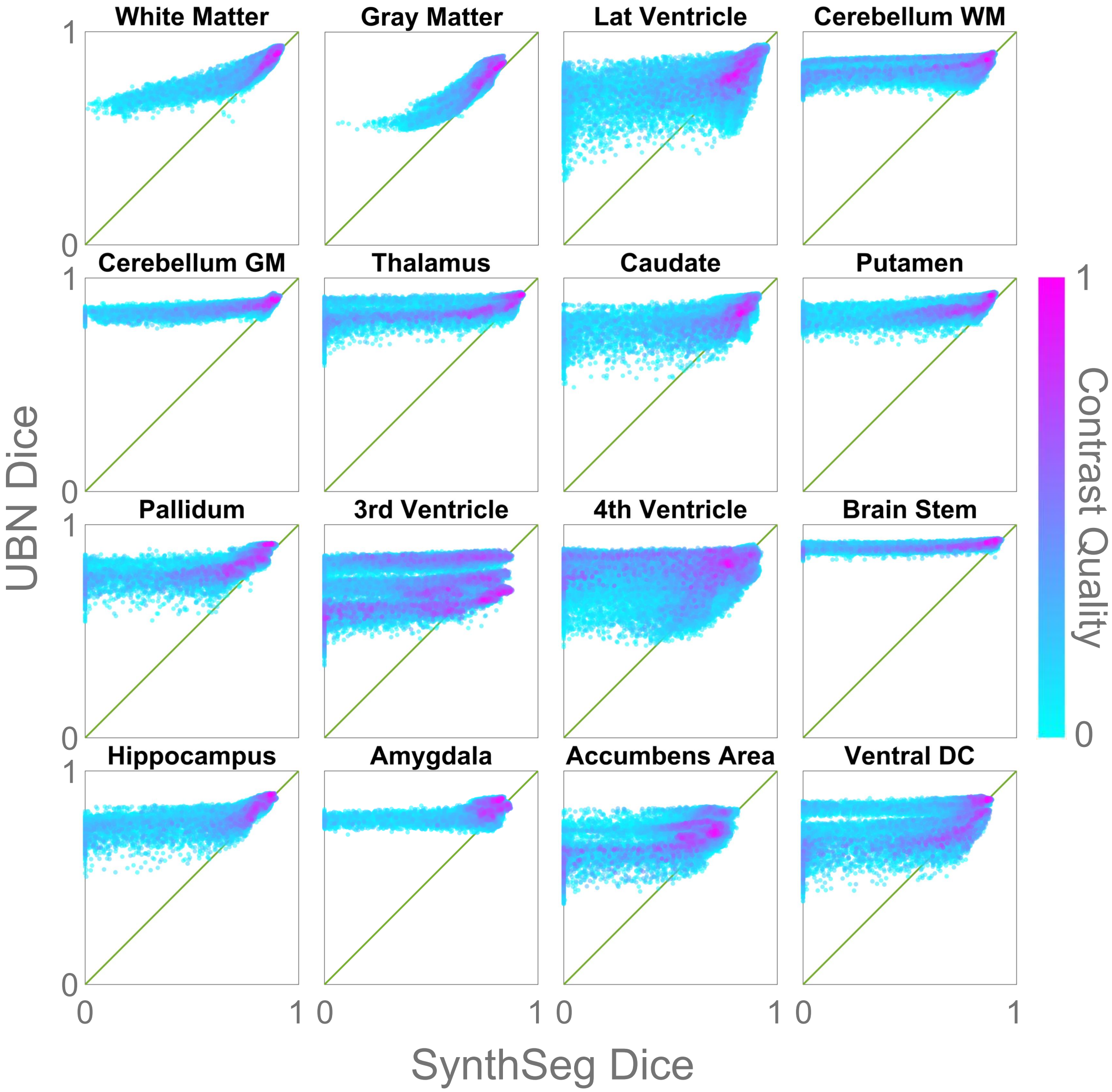


**Fig. S1** | Comparison of label-specific Dice scores for UBN and SynthSeg across 150,000 synthetic pan-contrast images. Y = X plotted in green for visual reference.

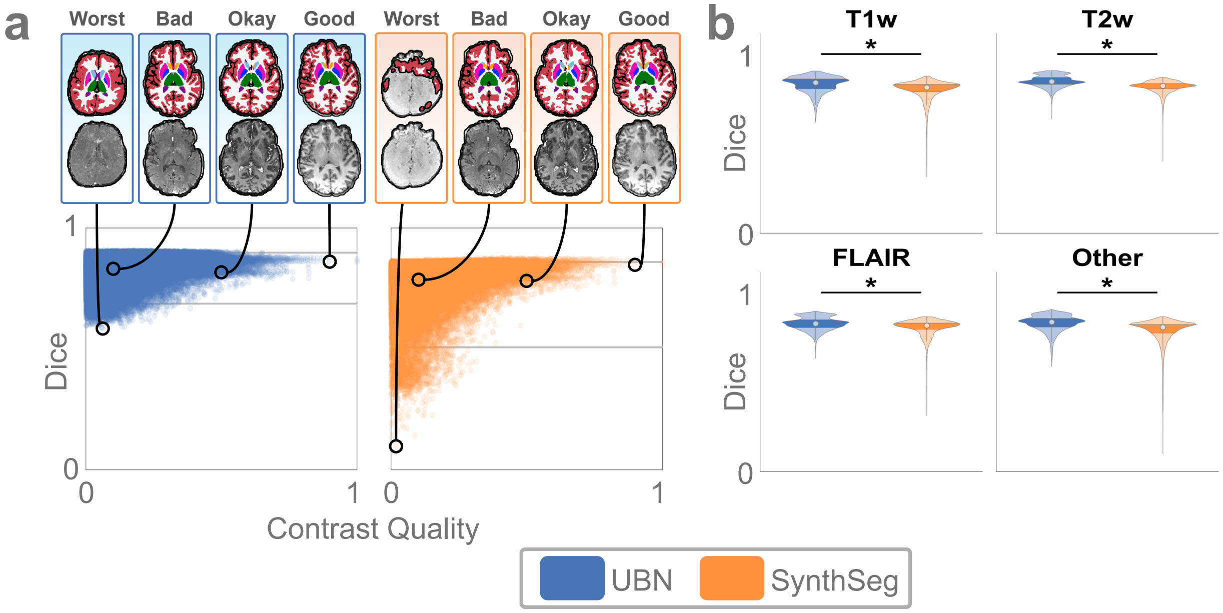


**Fig. S2** | Segmentation performance as a function of contrast quality and type. **a**, Whole-brain Dice scores plotted against contrast quality for 150,000 UltimateSynth images. Example UBN segmentations are shown for representative cases: 'Worst' corresponds to the lowest Dice score in the dataset; 'Bad', 'Okay', and 'Good' correspond to median Dice scores for images with contrast quality of $0.1 \pm 0.01$, $0.5 \pm 0.01$, and $0.9 \pm 0.01$, respectively. Gray lines indicate the 1st and 99th percentiles of Dice scores. **b**, Segmentation performance across scan types. The dataset was divided into 25,000 each of T1-weighted, T2-weighted, and FLAIR images, and 75,000 images with other contrasts. UBN significantly outperforms the SOTA in all four groups ($P < .05$).

**Tab. S2** | Dice scores by MR scan type for 150,000 UltimateSynth images show that UBN significantly outperforms SynthSeg across all four scan types ($P < .001$).

| MR Image Type | UBN Dice | SynthSeg Dice | Dice Difference *P*-Value |
|---|---|---|---|
| T1w | **.83 ± .05** | .80 ± .06 | **< .001** |
| T2w | **.85 ± .03** | .82 ± .03 | **< .001** |
| FLAIR | **.83 ± .04** | .80 ± .04 | **< .001** |
| Other | **.83 ± .05** | .78 ± .07 | **< .001** |

**Tab. S3** | Structure-specific Dice scores averaged across 160 T1-weighted and FLAIR scans from 10 ON-Harmony subjects (Figure 6b) show that UBN significantly outperforms both SOTA methods in 11 out of 16 labels ($P \leq .002$).

| Structure Label | UBN Dice | SynthSeg Dice | PhysSeg Dice | UBN-SS Dice Difference *P*-Value | UBN-PS Dice Difference *P*-Value | SS-PS Dice Difference *P*-Value |
|---|---|---|---|---|---|---|
| White Matter | **.93 ± .02** | .92 ± .01 | .87 ± .01 | **< .001** | **< .001** | **< .001** |
| Gray Matter | **.88 ± .02** | .86 ± .02 | .76 ± .02 | **< .001** | **< .001** | **< .001** |
| Lateral Ventricle | **.89 ± .03** | .88 ± .04 | - | **< .001** | - | - |
| Cerebellar White Matter | **.84 ± .02** | .83 ± .02 | - | **< .001** | - | - |
| Cerebellar Grey Matter | **.91 ± .01** | **.91 ± .01** | - | **< .001** | - | - |
| Thalamus | **.91 ± .01** | .89 ± .01 | - | **< .001** | - | - |
| Caudate | **.89 ± .02** | .88 ± .02 | - | **.002** | - | - |
| Putamen | **.90 ± .01** | .87 ± .02 | - | **< .001** | - | - |
| Pallidum | **.85 ± .02** | .77 ± .04 | - | **< .001** | - | - |
| 3rd Ventricle | .78 ± .05 | **.79 ± .07** | - | **.007** | - | - |
| 4th Ventricle | **.81 ± .04** | **.81 ± .05** | - | .16 | - | - |
| Brain Stem | **.93 ± .00** | .92 ± .01 | - | **< .001** | - | - |
| Hippocampus | **.88 ± .02** | .86 ± .02 | - | **< .001** | - | - |
| Amygdala | **.85 ± .02** | .84 ± .02 | - | .85 | - | - |
| Accumbens Area | **.74 ± .04** | .73 ± .03 | - | .21 | - | - |
| Ventral Diencephalon | **.85 ± .01** | .83 ± .02 | - | **< .001** | - | - |

**Tab. S4** | Structure-specific LVV, averaged across 160 T1-weighted and FLAIR scans from 10 ON-Harmony subjects (Figure 6b), shows that UBN significantly reduces LVV compared to SynthSeg and PhysSeg in 13 out of 16 structures ($P < .001$).

| Structure Label | UBN LVV [%] | SynthSeg LVV [%] | PhysSeg LVV [%] | UBN-SS LVV Difference *P*-Value | UBN-PS LVV Difference *P*-Value | SS-PS LVV Difference *P*-Value |
|---|---|---|---|---|---|---|
| White Matter | 2.92 ± 0.97 | 2.41 ± 1.29 | **2.27 ± 2.31** | **.002** | **< .001** | **< .001** |
| Gray Matter | **1.25 ± 1.00** | 2.35 ± 1.34 | 1.57 ± 1.53 | **< .001** | **< .001** | **< .001** |
| Lateral Ventricle | **1.70 ± 1.44** | 2.30 ± 1.82 | - | **< .001** | - | - |
| Cerebellar White Matter | **2.17 ± 1.16** | 9.24 ± 2.78 | - | **< .001** | - | - |
| Cerebellar Grey Matter | 3.46 ± 1.38 | **1.61 ± 1.30** | - | **< .001** | - | - |
| Thalamus | **0.84 ± 0.58** | 3.50 ± 2.34 | - | **< .001** | - | - |
| Caudate | **0.85 ± 0.68** | 1.41 ± 1.27 | - | **< .001** | - | - |
| Putamen | **2.61 ± 1.05** | 3.18 ± 2.54 | - | .07 | - | - |
| Pallidum | **1.28 ± 0.89** | 6.02 ± 4.39 | - | **< .001** | - | - |
| 3rd Ventricle | **4.71 ± 2.08** | 9.71 ± 6.00 | - | **< .001** | - | - |
| 4th Ventricle | **3.14 ± 2.08** | 15.65 ± 7.54 | - | **< .001** | - | - |
| Brain Stem | **1.25 ± 1.08** | 1.76 ± 1.35 | - | **< .001** | - | - |
| Hippocampus | **1.30 ± 1.00** | 3.69 ± 2.28 | - | **< .001** | - | - |
| Amygdala | **1.97 ± 1.34** | 6.51 ± 4.03 | - | **< .001** | - | - |
| Accumbens Area | **1.67 ± 1.26** | 4.11 ± 2.99 | - | **< .001** | - | - |
| Ventral Diencephalon | **2.18 ± 1.16** | 3.30 ± 2.47 | - | **< .001** | - | - |

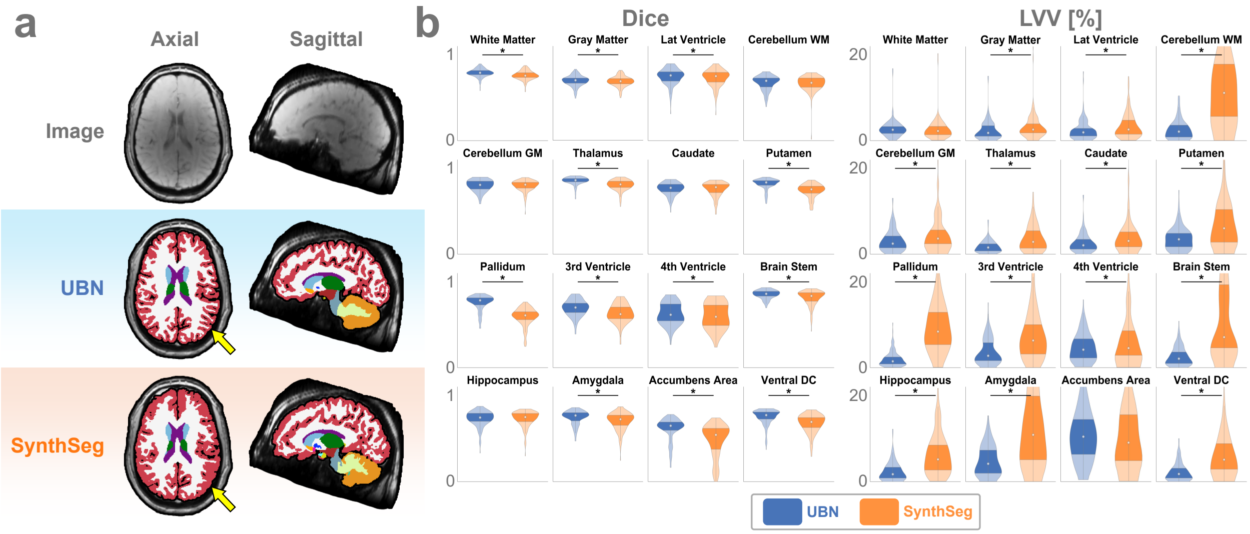


**Fig. S3** | SWI segmentation results across 160 ON-Harmony scans. **a**, UBN preserves cortical details even in the inherently low-contrast SWI scans (see yellow arrows). **b**, Quantitative comparison of Dice and LVV scores for UBN and SynthSeg. UBN achieves significantly higher Dice scores in 12 of 16 labels and significantly lower LVV in 14 of 16 labels.

**Tab. S5** | Structure-specific Dice and LVV scores across 160 SWI scans from 10 ON-Harmony subjects (Figure S3). Despite low image contrast and reduced through-plane resolution, UBN achieves significantly higher Dice in 11 structures ($P < .001$) and significantly lower LVV in 14 structures ($P \leq .006$).

| Structure Label | UBN Dice | SynthSeg Dice | Dice Difference *P*-Value | UBN LVV [%] | SynthSeg LVV [%] | LVV Difference *P*-Value |
|---|---|---|---|---|---|---|
| White Matter | **.79 ± .05** | .76 ± .05 | **< .001** | 2.72 ± 2.02 | **2.56 ± 2.38** | .06 |
| Gray Matter | **.69 ± .06** | **.69 ± .06** | **< .001** | **2.33 ± 2.16** | 3.16 ± 2.75 | **< .001** |
| Lateral Ventricle | **.75 ± .10** | .73 ± .10 | **< .001** | **2.42 ± 2.23** | 3.38 ± 2.86 | **< .001** |
| Cerebellar White Matter | **.67 ± .08** | **.67 ± .09** | .67 | **2.54 ± 2.20** | 13.21 ± 11.92 | **< .001** |
| Cerebellar Grey Matter | **.80 ± .07** | **.80 ± .06** | .21 | **3.08 ± 2.45** | 5.34 ± 6.67 | **< .001** |
| Thalamus | **.85 ± .05** | .80 ± .06 | **< .001** | **1.73 ± 1.37** | 3.65 ± 2.96 | **< .001** |
| Caudate | **.76 ± .07** | **.76 ± .08** | .22 | **2.55 ± 2.14** | 4.19 ± 4.47 | **< .001** |
| Putamen | **.82 ± .06** | .74 ± .08 | **< .001** | **3.56 ± 2.55** | 7.44 ± 6.65 | **< .001** |
| Pallidum | **.76 ± .09** | .59 ± .10 | **< .001** | **1.96 ± 1.72** | 9.46 ± 6.04 | **< .001** |
| 3rd Ventricle | **.70 ± .09** | .63 ± .11 | **< .001** | **3.74 ± 2.98** | 7.82 ± 6.57 | **< .001** |
| 4th Ventricle | **.63 ± .12** | .60 ± .15 | **< .001** | **4.68 ± 3.20** | 6.87 ± 9.12 | **.006** |
| Brain Stem | **.85 ± .05** | .81 ± .09 | **< .001** | **2.62 ± 2.14** | 12.86 ± 12.23 | **< .001** |
| Hippocampus | **.73 ± .09** | **.73 ± .09** | .56 | **2.35 ± 2.09** | 6.19 ± 4.62 | **< .001** |
| Amygdala | **.74 ± .08** | .70 ± .10 | **< .001** | **4.83 ± 3.40** | 13.69 ± 11.39 | **< .001** |
| Accumbens Area | **.61 ± .13** | .48 ± .19 | **< .001** | **11.16 ± 6.68** | 12.16 ± 12.06 | .97 |
| Ventral Diencephalon | **.74 ± .09** | .67 ± .11 | **< .001** | **2.24 ± 2.02** | 6.63 ± 7.27 | **< .001** |

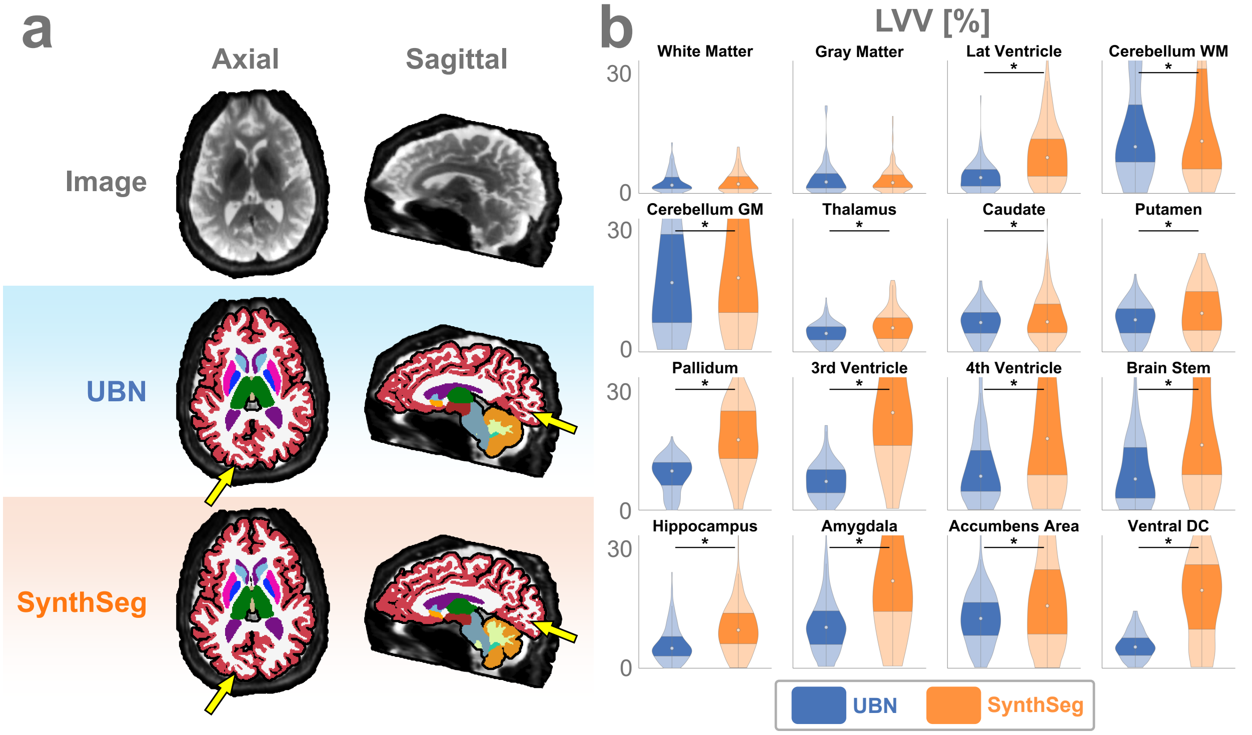


**Fig. S4** | DWI segmentation examples and quantitative results across 160 ON-Harmony images. **a**, UBN is robust to spatial inhomogeneity and poor contrast, particularly in the occipital lobe (yellow arrows). **b**, LVV scores for UBN and SynthSeg on the 160 DWI test images. Despite mild performance drops in cerebral WM and GM due to DWI-related distortions, UBN still achieves strong overall performance, with significant improvements over the SOTA in 13 of 16 labels.

**Tab. S6** | Structure-specific LVV results from 160 DWI images across 10 subjects in the ON-Harmony dataset (Figure S4). Despite pronounced geometric distortions, UBN consistently produces significantly lower LVV than SynthSeg in 14 out of 16 structure labels ($P \leq .03$).

| Structure Label | UBN LVV [%] | SynthSeg LVV [%] | LVV Difference *P*-Value |
|---|---|---|---|
| White Matter | **2.97 ± 2.56** | 3.03 ± 2.60 | .75 |
| Gray Matter | 3.75 ± 3.82 | **3.49 ± 3.02** | .92 |
| Lateral Ventricle | **4.24 ± 3.29** | 10.00 ± 7.37 | **< .001** |
| Cerebellar White Matter | **16.27 ± 13.89** | 22.40 ± 22.54 | **.005** |
| Cerebellar Grey Matter | **19.75 ± 15.04** | 24.40 ± 19.75 | **.006** |
| Thalamus | **4.61 ± 2.52** | 6.35 ± 4.03 | **< .001** |
| Caudate | **7.16 ± 3.84** | 8.51 ± 5.48 | **.03** |
| Putamen | **7.81 ± 4.18** | 10.13 ± 6.26 | **< .001** |
| Pallidum | **8.73 ± 4.17** | 18.11 ± 8.70 | **< .001** |
| 3rd Ventricle | **7.36 ± 4.43** | 28.06 ± 17.17 | **< .001** |
| 4th Ventricle | **11.84 ± 11.85** | 25.56 ± 25.61 | **< .001** |
| Brain Stem | **9.96 ± 8.75** | 22.90 ± 20.71 | **< .001** |
| Hippocampus | **5.88 ± 4.33** | 10.18 ± 7.41 | **< .001** |
| Amygdala | **11.22 ± 7.83** | 27.78 ± 19.91 | **< .001** |
| Accumbens Area | **12.40 ± 6.63** | 17.56 ± 12.53 | **< .001** |
| Ventral Diencephalon | **5.65 ± 3.29** | 18.85 ± 12.02 | **< .001** |

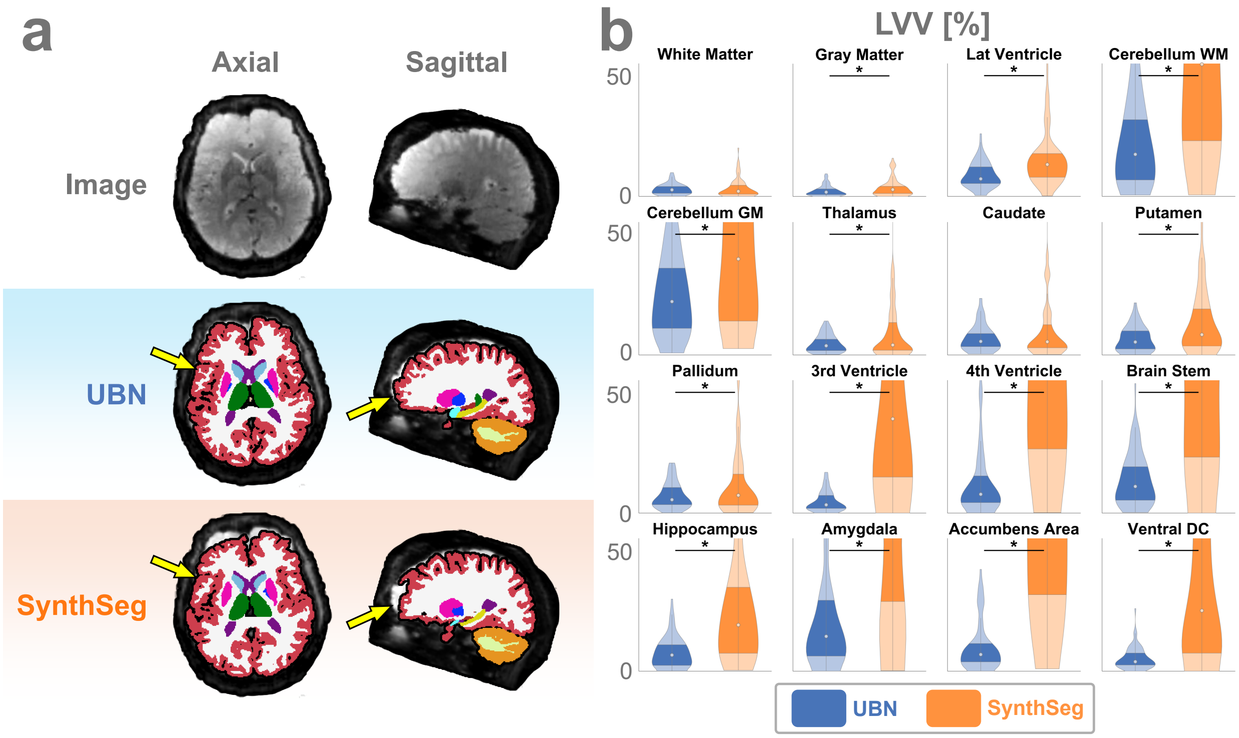


**Fig. S5** | Resting-state fMRI segmentation examples and quantitative results across 80 ON-Harmony images. **a**, UBN generates anatomically plausible segmentations that are less affected by low SNR and spatial inhomogeneity compared to SynthSeg (yellow arrows). **b**, LVV scores for UBN and SynthSeg on the 80 fMRI test images. While poor image quality increases variation for both methods, SynthSeg's errors—particularly in small basal ganglia structures—result in substantially higher LVV. UBN achieves significantly lower LVV in 14 out of 16 labels.

**Tab. S7** | Structure-specific LVV across 80 resting-state fMRI images from 10 ON-Harmony subjects (Figure S5). Low resolution and signal dropout slightly degrade LVV for both UBN and SOTA methods. Despite this, UBN's segmentations show significantly less volume variation from the cross-image mean than SynthSeg's in 14 structures ($P \leq .03$).

| Structure Label | UBN LVV [%] | SynthSeg LVV [%] | LVV Difference *P*-Value |
|---|---|---|---|
| White Matter | **2.97 ± 2.25** | 3.44 ± 3.83 | .81 |
| Gray Matter | **2.37 ± 2.19** | 3.62 ± 3.71 | **.01** |
| Lateral Ventricle | **8.79 ± 5.14** | 15.36 ± 11.26 | **< .001** |
| Cerebellar White Matter | **21.52 ± 17.37** | 57.92 ± 39.42 | **< .001** |
| Cerebellar Grey Matter | **24.96 ± 16.75** | 42.04 ± 28.99 | **< .001** |
| Thalamus | **4.66 ± 3.62** | 9.53 ± 12.16 | **.02** |
| Caudate | 7.00 ± 5.34 | 10.08 ± 12.08 | .46 |
| Putamen | **6.55 ± 5.08** | 13.32 ± 12.72 | **< .001** |
| Pallidum | **7.24 ± 5.43** | 11.16 ± 12.26 | **.03** |
| 3rd Ventricle | **4.82 ± 4.16** | 57.63 ± 62.60 | **< .001** |
| 4th Ventricle | **13.61 ± 15.75** | 67.86 ± 54.58 | **< .001** |
| Brain Stem | **13.58 ± 10.57** | 63.77 ± 46.95 | **< .001** |
| Hippocampus | **7.68 ± 6.27** | 25.21 ± 21.65 | **< .001** |
| Amygdala | **23.58 ± 26.20** | 83.36 ± 95.96 | **< .001** |
| Accumbens Area | **10.19 ± 10.06** | 72.73 ± 62.67 | **< .001** |
| Ventral Diencephalon | **5.25 ± 4.41** | 36.91 ± 37.12 | **< .001** |

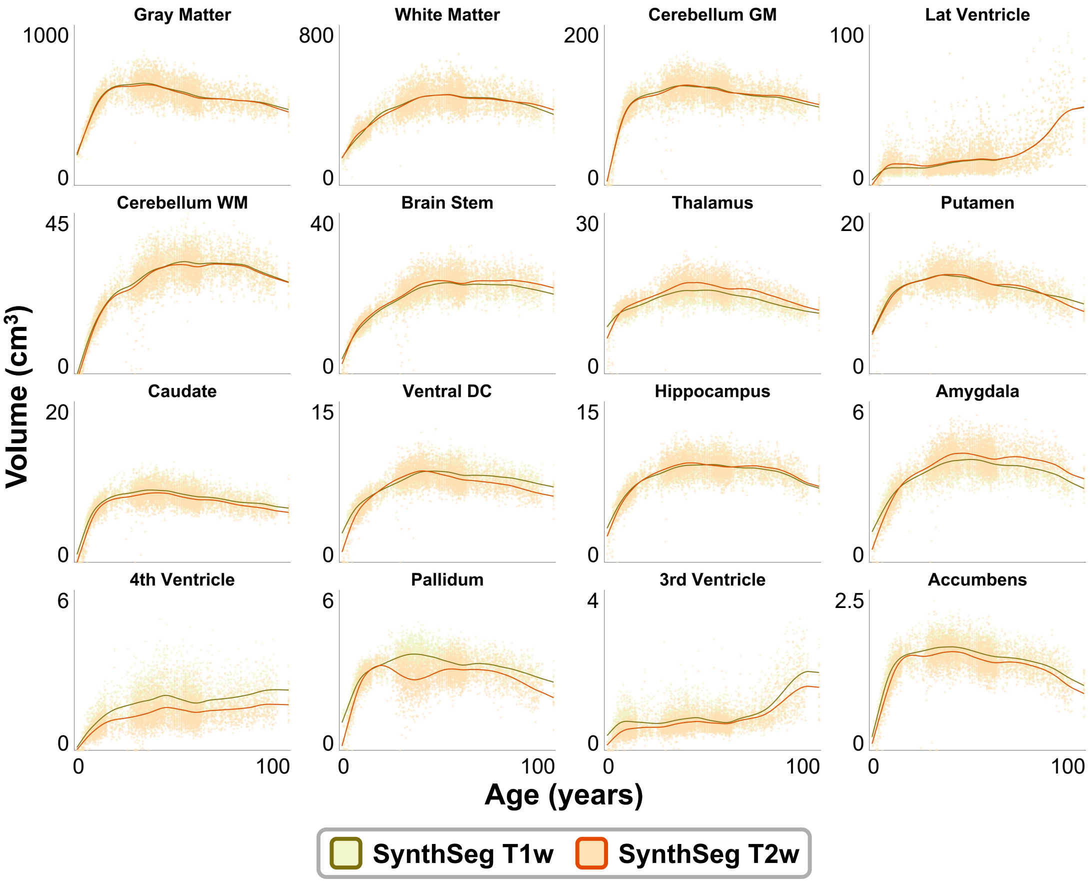


**Fig. S6** | Label volumes from SynthSeg segmentations of 8,385 T1w and T2w images spanning the human lifespan. Volume trends are shown separately for T1w (yellow) and T2w (orange) scans, revealing notable intra-method discrepancies in typical volumes for several structures, especially small deep brain regions such as the thalamus, pallidum, and accumbens area.

**Tab. S8** | UBN volume mean absolute errors (MAEs) on 4,090 paired T1w and T2w lifespan MR images (Figure 7). UBN significantly outperforms the SOTA in 12 labels.

| Structure Label | UBN MAE [%] | SynthSeg MAE [%] | MAE Difference *P*-Value |
|---|---|---|---|
| White Matter | 3.20 ± 2.05 | **2.37 ± 3.71** | **< .001** |
| Gray Matter | 2.31 ± 1.71 | **1.81 ± 2.28** | **< .001** |
| Lateral Ventricle | **8.30 ± 3.53** | 8.70 ± 13.00 | **< .001** |
| Cerebellar White Matter | **1.82 ± 1.76** | 5.80 ± 13.83 | **< .001** |
| Cerebellar Grey Matter | 3.18 ± 1.98 | **3.13 ± 9.72** | **< .001** |
| Thalamus | **2.67 ± 1.36** | 7.31 ± 7.01 | **< .001** |
| Caudate | **4.47 ± 4.11** | 6.60 ± 12.96 | **< .001** |
| Putamen | 4.36 ± 1.74 | **3.17 ± 5.27** | **< .001** |
| Pallidum | **3.65 ± 2.62** | 17.17 ± 17.08 | **< .001** |
| 3rd Ventricle | **5.28 ± 4.42** | 20.10 ± 19.98 | **< .001** |
| 4th Ventricle | **4.00 ± 3.78** | 27.59 ± 18.60 | **< .001** |
| Brain Stem | **1.61 ± 1.47** | 5.59 ± 11.80 | **< .001** |
| Hippocampus | **2.71 ± 1.91** | 3.61 ± 5.55 | **< .001** |
| Amygdala | **3.04 ± 2.27** | 8.35 ± 10.73 | **< .001** |
| Accumbens Area | **2.87 ± 2.48** | 8.63 ± 11.69 | **< .001** |
| Ventral Diencephalon | **2.25 ± 2.00** | 7.84 ± 13.43 | **< .001** |

**Tab. S9** | UBN performance on 2,030 low-resolution, low-field M4Raw images (Figure 9). UBN significantly reduces LVV compared to the SOTA across 15 labels.

| Structure Label | UBN LVV [%] | SynthSeg LVV [%] | LVV Difference *P*-Value |
|---|---|---|---|
| White Matter | **3.40 ± 2.61** | 4.45 ± 4.26 | **< .001** |
| Gray Matter | 2.63 ± 2.11 | **1.71 ± 1.31** | **< .001** |
| Lateral Ventricle | **4.81 ± 3.24** | 7.49 ± 5.47 | **< .001** |
| Cerebellar White Matter | **8.66 ± 12.39** | 11.08 ± 11.04 | **< .001** |
| Cerebellar Grey Matter | **3.41 ± 4.29** | 5.91 ± 5.96 | **< .001** |
| Thalamus | **1.96 ± 1.77** | 4.64 ± 3.66 | **< .001** |
| Caudate | **4.13 ± 4.66** | 8.67 ± 8.90 | **< .001** |
| Putamen | **2.04 ± 1.61** | 5.76 ± 5.24 | **< .001** |
| Pallidum | **2.57 ± 2.10** | 9.64 ± 7.43 | **< .001** |
| 3rd Ventricle | **6.80 ± 4.55** | 8.68 ± 6.48 | **< .001** |
| 4th Ventricle | **6.61 ± 6.89** | 15.74 ± 12.00 | **< .001** |
| Brain Stem | **1.75 ± 1.53** | 3.08 ± 3.03 | **< .001** |
| Hippocampus | **2.16 ± 2.09** | 3.89 ± 2.93 | **< .001** |
| Amygdala | **2.86 ± 2.52** | 7.32 ± 5.89 | **< .001** |
| Accumbens Area | **4.10 ± 3.36** | 7.88 ± 6.26 | **< .001** |
| Ventral Diencephalon | **2.81 ± 2.24** | 3.86 ± 2.98 | **< .001** |

**Tab. S10** | Structure-specific mean absolute error across 8,180 paired T1w and T2w MR images (Figure 10). UCN shows notably low volume error in both cerebellar labels. CerebNet is excluded due to incompatibility with T2w images.

| Structure Label | UCN MAE [%] |
|---|---|
| Cerebellar White Matter | 2.69 ± 2.13 |
| Cerebellar Gray Matter | 2.20 ± 2.38 |

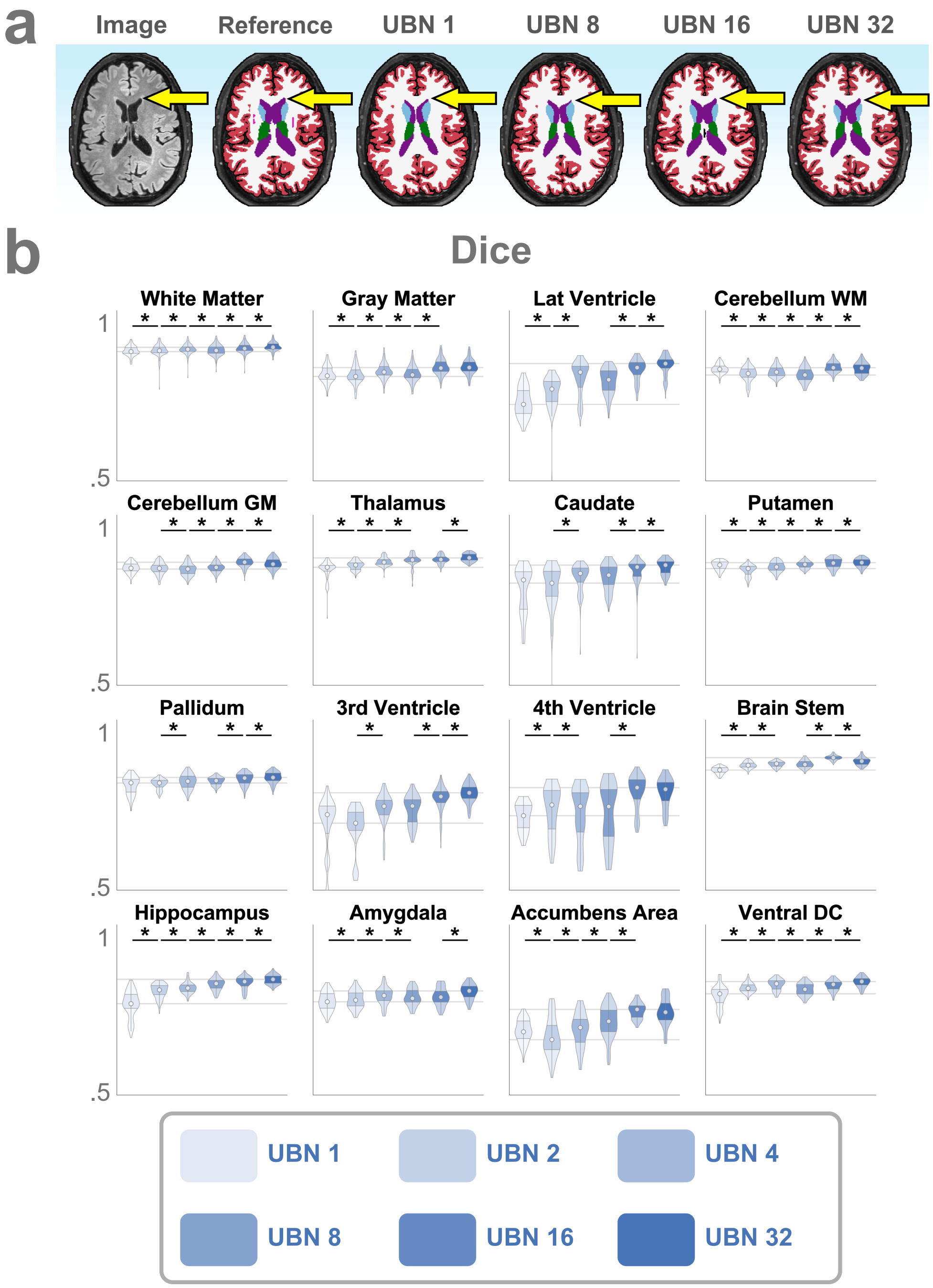


**Fig. S7** | UBN performance versus number of unique training anatomies. **a**, Compared to FreeSurfer consensus labels, UBN trained on a single anatomy (UBN 1) sometimes missegments small structures like the lateral ventricle (yellow arrows), while training on 32 diverse anatomies (UBN 32) improves segmentation accuracy. **b**, Dice scores across networks trained with varying numbers of unique anatomies, evaluated on 160 T1w and FLAIR test images against FreeSurfer references. Dice performance generally improves with larger training sets, with 9 of 16 labels showing significant gains between 16 and 32 training anatomies ($P < .01$). Gray lines indicate the minimum and maximum median Dice scores across all networks to highlight these improvements.

**Tab. S11** | UBN performance versus training sample size (Figure S7), measured by structure-specific Dice scores averaged over 160 T1-weighted and FLAIR scans from 10 ON-Harmony subjects. Dice scores improve as the number of training anatomies increases but plateau between UBN 16 and UBN 32.

| Structure Label | UBN 1 Dice | UBN 2 Dice | UBN 4 Dice | UBN 8 Dice | UBN 16 Dice | UBN 32 Dice | 1-2 Dice Difference *P*-Value | 2-4 Dice Difference *P*-Value | 4-8 Dice Difference *P*-Value | 8-16 Dice Difference *P*-Value | 16-32 Dice Difference *P*-Value |
|---|---|---|---|---|---|---|---|---|---|---|---|
| White Matter | .92 ± .02 | .92 ± .02 | **.93 ± .02** | .92 ± .02 | **.93 ± .02** | **.93 ± .01** | **< .001** | **< .001** | **< .001** | **< .001** | **< .001** |
| Gray Matter | .84 ± .03 | .84 ± .03 | .85 ± .02 | .85 ± .03 | **.87 ± .03** | **.87 ± .02** | **.01** | **< .001** | **< .001** | **< .001** | .48 |
| Lateral Ventricle | .75 ± .04 | .79 ± .05 | .83 ± .06 | .82 ± .04 | .86 ± .03 | **.87 ± .03** | **< .001** | **< .001** | **.02** | **< .001** | **< .001** |
| Cerebellar White Matter | .86 ± .02 | .84 ± .02 | .85 ± .02 | .84 ± .02 | **.87 ± .02** | .86 ± .02 | **< .001** | **< .001** | **< .001** | **< .001** | **< .001** |
| Cerebellar Grey Matter | .88 ± .02 | .88 ± .02 | .87 ± .02 | .88 ± .02 | **.90 ± .02** | .89 ± .02 | .11 | **< .001** | **< .001** | **< .001** | **< .001** |
| Thalamus | .87 ± .03 | .89 ± .01 | .90 ± .01 | .90 ± .02 | **.91 ± .01** | **.91 ± .01** | **< .001** | **< .001** | **< .001** | .08 | **< .001** |
| Caudate | .81 ± .07 | .82 ± .07 | .85 ± .04 | .85 ± .04 | .87 ± .04 | **.88 ± .02** | .02 | **< .001** | .43 | **< .001** | **< .001** |
| Putamen | .89 ± .01 | .87 ± .02 | .88 ± .02 | .89 ± .01 | .89 ± .02 | **.90 ± .01** | **< .001** | **< .001** | **< .001** | **< .001** | **.002** |
| Pallidum | .84 ± .03 | .84 ± .02 | .85 ± .02 | .85 ± .01 | **.86 ± .02** | **.86 ± .02** | .04 | **< .001** | .71 | **< .001** | **< .001** |
| 3rd Ventricle | .71 ± .09 | .70 ± .07 | .76 ± .05 | .76 ± .05 | .80 ± .04 | **.81 ± .03** | .03 | **< .001** | .02 | **< .001** | **< .001** |
| 4th Ventricle | .74 ± .04 | .76 ± .08 | .74 ± .09 | .75 ± .09 | **.82 ± .05** | **.82 ± .05** | **< .001** | **< .001** | .46 | **< .001** | .90 |
| Brain Stem | .89 ± .01 | .90 ± .01 | .91 ± .01 | .91 ± .01 | **.93 ± .01** | .92 ± .01 | **< .001** | **< .001** | .83 | **< .001** | **< .001** |
| Hippocampus | .79 ± .04 | .84 ± .02 | .85 ± .02 | .86 ± .02 | .86 ± .02 | **.87 ± .01** | **< .001** | **< .001** | **< .001** | **< .001** | **< .001** |
| Amygdala | .80 ± .03 | .81 ± .03 | .82 ± .03 | .82 ± .03 | .82 ± .03 | **.83 ± .02** | **< .001** | **< .001** | **.001** | .15 | **< .001** |
| Accumbens Area | .71 ± .04 | .69 ± .06 | .71 ± .05 | .74 ± .05 | **.77 ± .02** | **.77 ± .04** | **< .001** | **< .001** | **< .001** | **< .001** | .43 |
| Ventral Diencephalon | .82 ± .03 | .85 ± .01 | **.86 ± .02** | .84 ± .02 | .85 ± .02 | **.86 ± .02** | **< .001** | **< .001** | **< .001** | **< .001** | **< .001** |

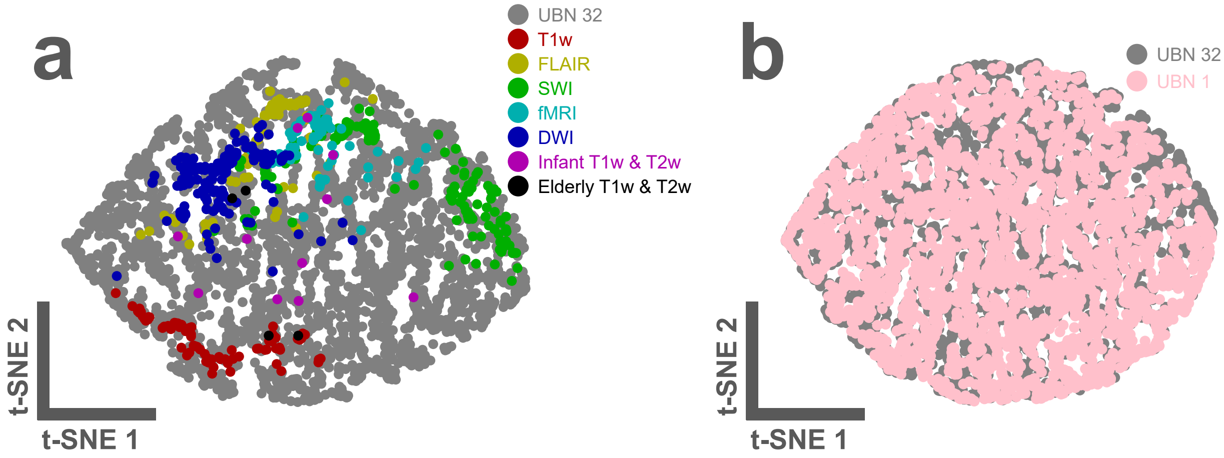


**Fig. S8** | t-SNE visualization of pan-contrast coverage in the UBN 32 and UBN 1 training datasets relative to a variety of test images. For each label, 15 first-order radiomic features were extracted, totaling 240 features per image. Each point in the 2D scatter plot represents one image. **a**, Comparison between UBN 32 training images and 575 external test images. Points cluster by contrast similarity, showing that the 3,200 UltimateSynth images in UBN 32's training data span the full range of common and uncommon test-time contrasts. T1-weighted, FLAIR, and atypical contrasts from the test set are well represented within the gray training point cloud. **b**, Overlap in image contrast distributions between UBN 1 and UBN 32 suggests that broad contrast diversity can be achieved independently of anatomical diversity.

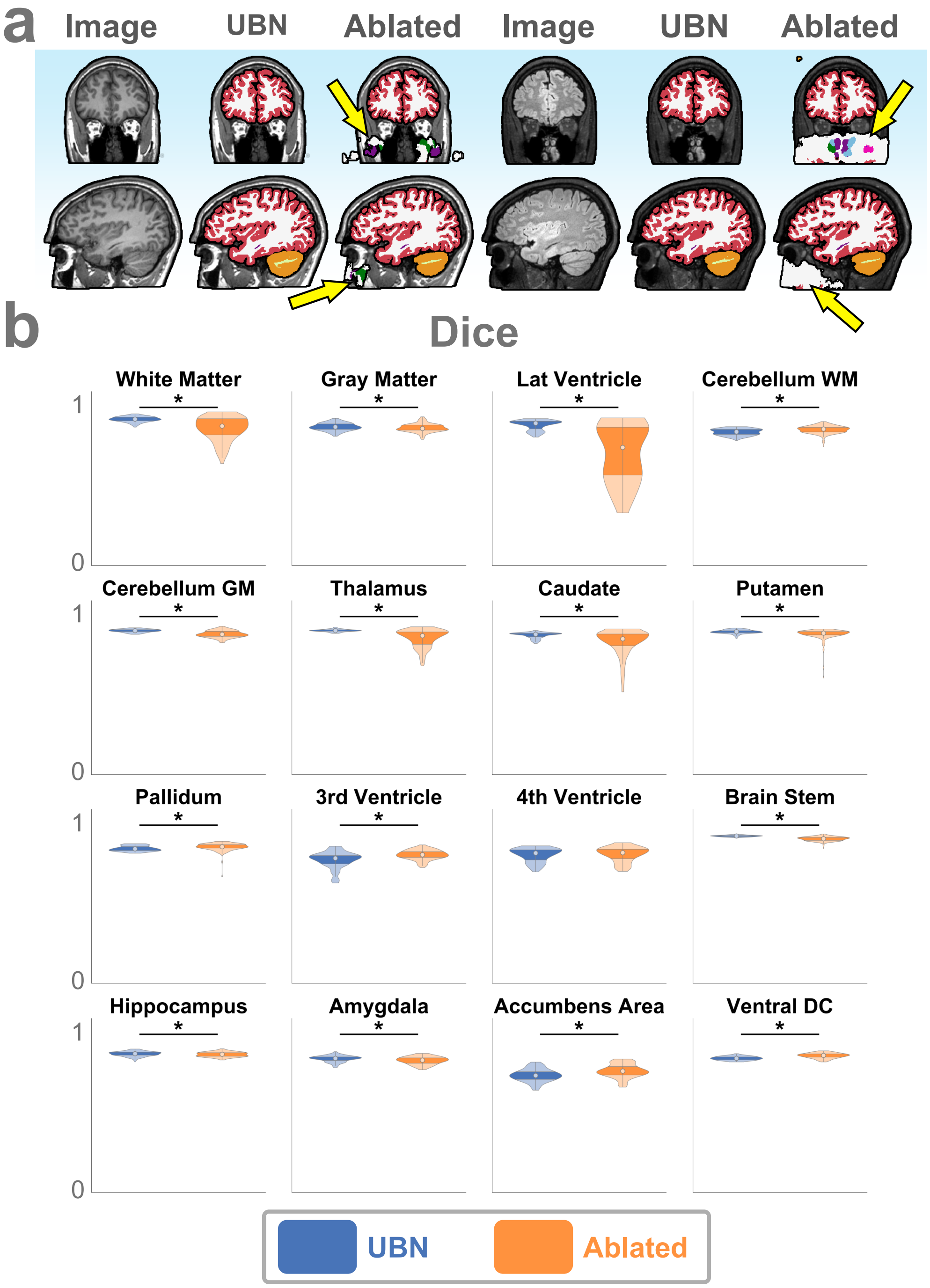


**Fig. S9** | Comparison of UBN and a data augmentation–ablated variant. **a**, Networks trained without background noise corruption or non-brain soft tissue simulation frequently mislabel regions outside the brain (yellow arrows). **b**, Dice scores show comparable performance in deep structures, but the ablated network struggles with larger regions like white matter and lateral ventricles. Across 160 T1w and FLAIR test images, UBN significantly outperforms the ablated variant in 10 of 16 labels relative to FreeSurfer reference labels ($P < .05$).

**Tab. S12** | UBN performance with and without extensive training data augmentation (Figure S9), evaluated using structure-specific Dice scores across 160 T1-weighted and FLAIR scans from 10 ON-Harmony subjects. UBN significantly outperforms the ablated model in 10 labels, with large improvements in white matter, lateral ventricles, and the caudate nucleus.

| Structure Label | UBN Dice | Ablated Dice | Dice Difference *P*-Value |
|---|---|---|---|
| White Matter | **.93 ± .02** | .87 ± .07 | **< .001** |
| Gray Matter | **.88 ± .02** | .87 ± .03 | **.01** |
| Lateral Ventricle | **.89 ± .03** | .71 ± .18 | **< .001** |
| Cerebellar White Matter | .84 ± .02 | **.86 ± .02** | **< .001** |
| Cerebellar Grey Matter | **.91 ± .01** | .89 ± .02 | **< .001** |
| Thalamus | **.91 ± .01** | .86 ± .06 | **< .001** |
| Caudate | **.88 ± .02** | .83 ± .08 | **< .001** |
| Putamen | **.90 ± .01** | .89 ± .04 | **< .001** |
| Pallidum | .85 ± .02 | **.86 ± .02** | **< .001** |
| 3rd Ventricle | .78 ± .05 | **.81 ± .03** | **< .001** |
| 4th Ventricle | .81 ± .04 | **.82 ± .05** | .06 |
| Brain Stem | **.93 ± .00** | .91 ± .01 | **< .001** |
| Hippocampus | **.88 ± .02** | .87 ± .01 | **.05** |
| Amygdala | **.85 ± .02** | .83 ± .02 | **< .001** |
| Accumbens Area | .74 ± .04 | **.77 ± .04** | **< .001** |
| Ventral Diencephalon | .85 ± .01 | **.86 ± .02** | **< .001** |

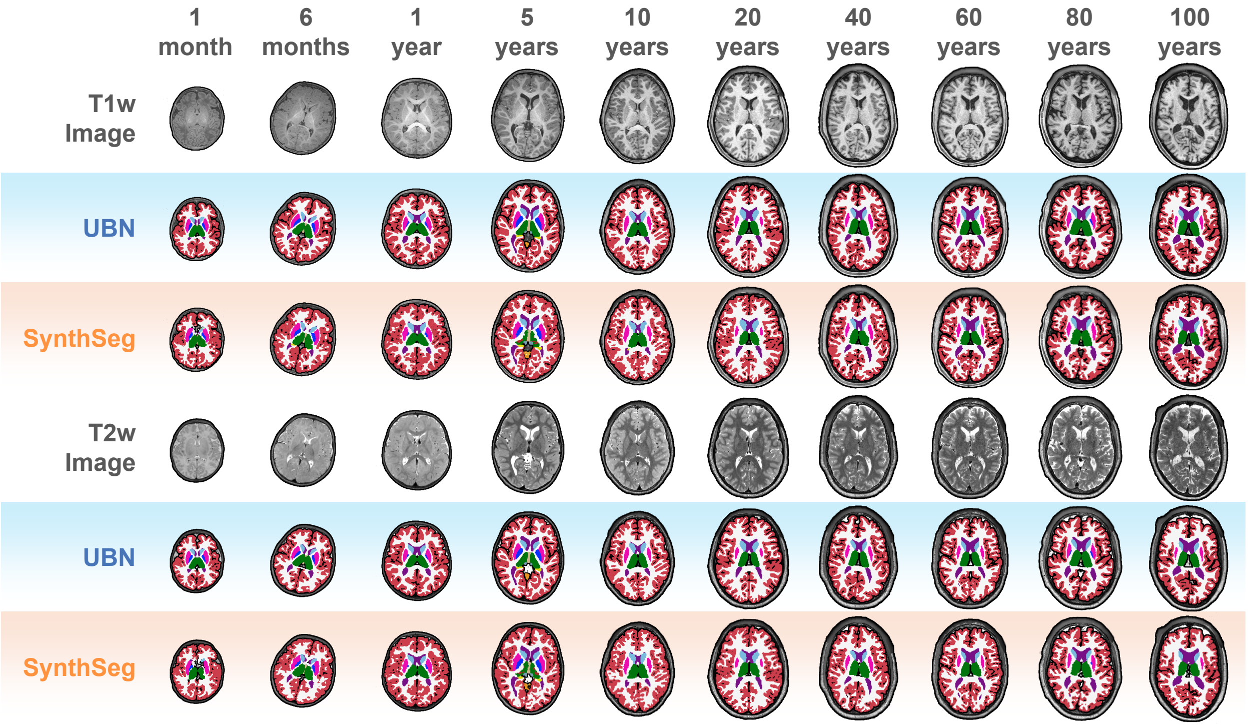


**Fig. S10** | Example segmentations from ten representative subjects across the human lifespan using UBN and SynthSeg in axial plane view.

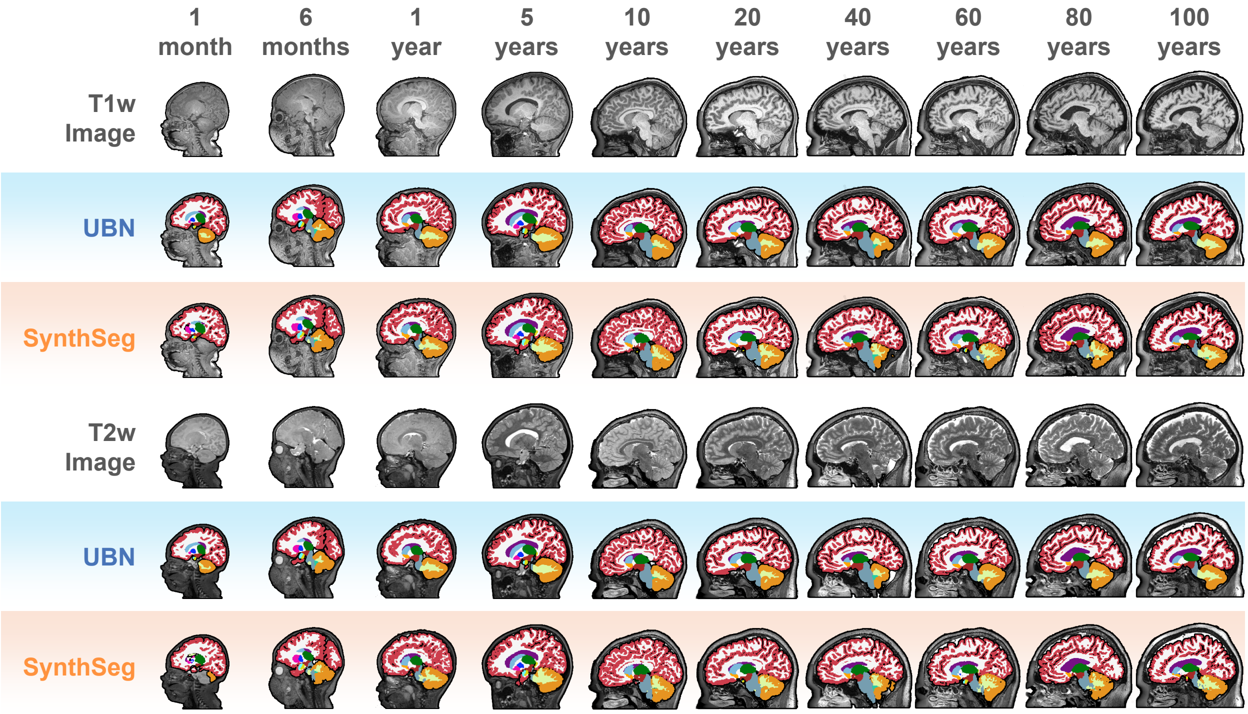


**Fig. S11** | Example segmentations from ten representative subjects across the human lifespan using UBN and SynthSeg in sagittal view.

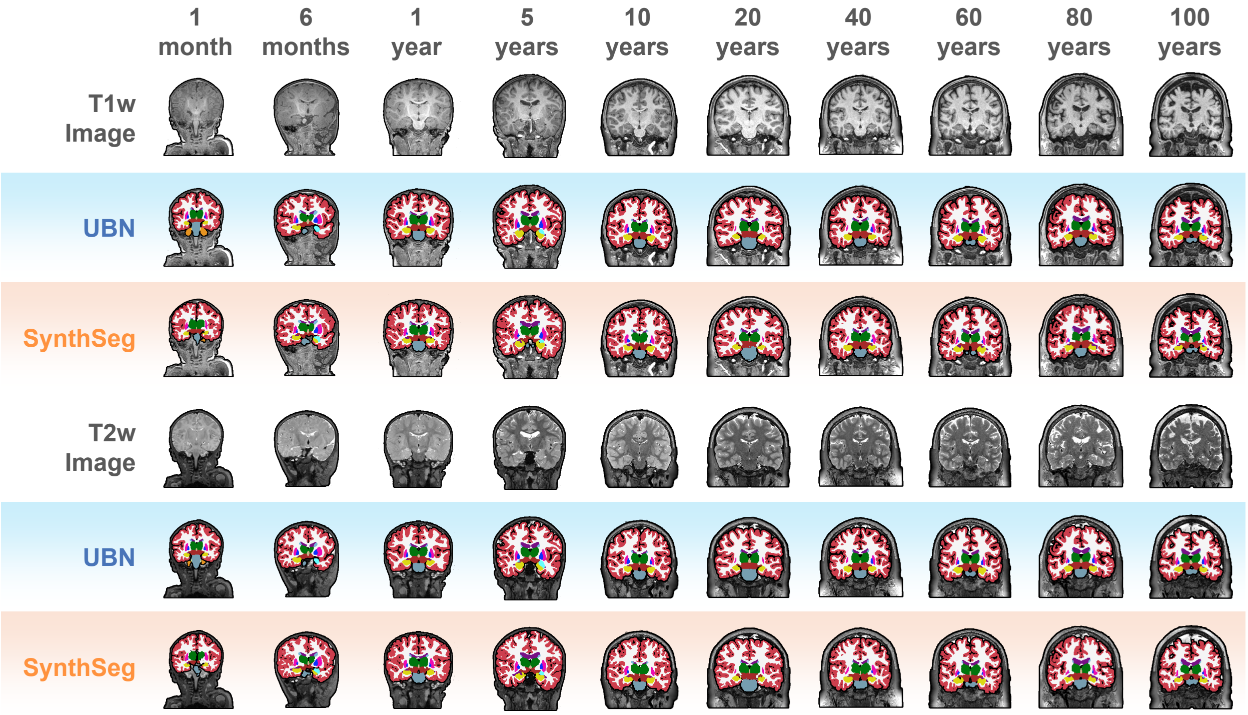


**Fig. S12** | Example segmentations from ten representative subjects across the human lifespan using UBN and SynthSeg in coronal view.

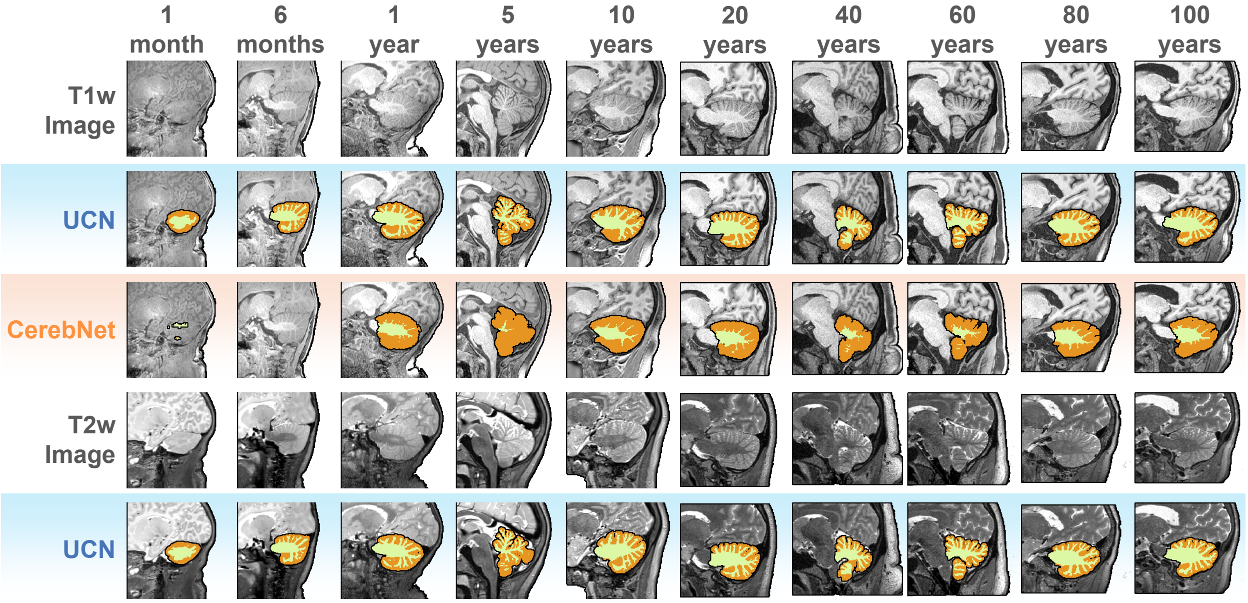


**Fig. S13** | Example segmentations from ten representative subjects across the human lifespan using UCN and CerebNet in sagittal view.

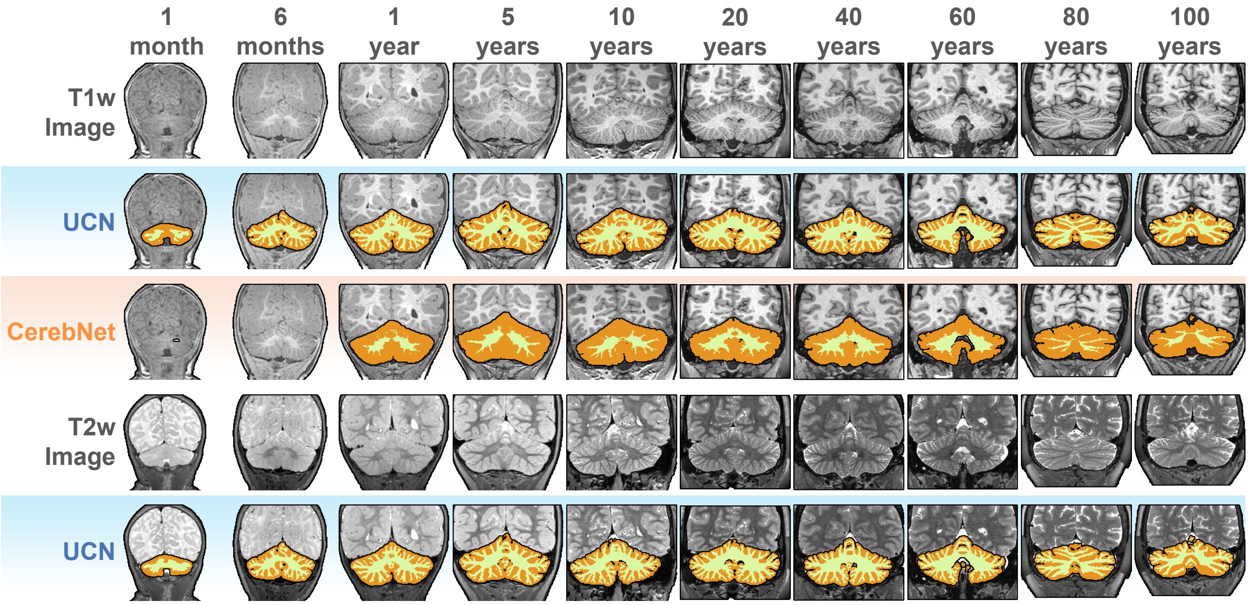


**Fig. S14** | Example segmentations from ten representative subjects across the human lifespan using UCN and CerebNet in coronal view.

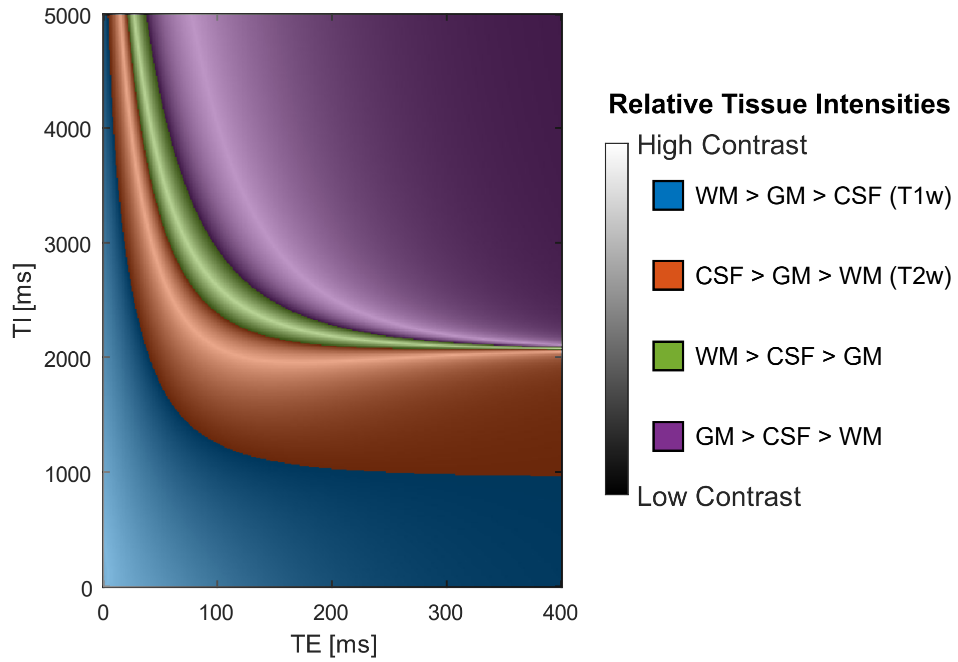


**Fig. S15** | Distribution of inter-tissue contrasts among gray matter, white matter, and cerebrospinal fluid across the scan parameter space of the UltimateSynth dictionary, shown here using TI and TE axes. Each colored region represents a unique ordering of tissue intensities, corresponding to familiar contrasts such as T1-weighted or T2-weighted contrasts. Bright bands indicate scan parameter combinations that yield high tissue contrast. Due to the exponential decay of magnetization over time, these high-contrast regions are unevenly distributed across the parameter space.
